# Constraints on the Emergence of RNA through Non-Templated Primer Extension with Mixtures of Potentially Prebiotic Nucleotides

**DOI:** 10.1101/2024.01.21.576316

**Authors:** Xiwen Jia, Stephanie J. Zhang, Lijun Zhou, Jack W. Szostak

## Abstract

The emergence of RNA on the early Earth is likely to have been influenced by a series of chemical and physical processes that acted to filter out various alternative nucleic acids. For example, UV photostability is thought to have favored the survival of the canonical nucleotides. In a recent proposal for the prebiotic synthesis of the building blocks of RNA, ribonucleotides share a common pathway with arabino- and threo-nucleotides. We have therefore investigated non-templated primer extension with 2-aminoimidazole-activated forms of these alternative nucleotides to see if the synthesis of the first oligonucleotides might have been biased in favor of RNA. We show that non-templated primer extension occurs predominantly through 5ʹ-5ʹ imidazolium bridged dinucleotides, echoing the mechanism of template-directed primer extension. Ribo- and arabino-nucleotides exhibited comparable rates and yields of non-templated primer extension, whereas threo-nucleotides showed lower reactivity. Competition experiments with mixtures of nucleotides confirmed the bias against the incorporation of threo-nucleotides into oligonucleotides. This bias, coupled with selective prebiotic synthesis and templated copying favoring ribonucleotides, provides a plausible model for the exclusion of threo-nucleotides from primordial oligonucleotides. In contrast, the exclusion of arabino-nucleotides may have resulted primarily from biases in synthesis and in template-directed primer extension.

## INTRODUCTION

RNA is considered to be a promising candidate for the primordial genetic polymer, owing to its dual roles in encoding genetic information and catalyzing reactions (1–3). However, just how RNA might have emerged from prebiotic mixtures remains an open question. Potentially prebiotic synthetic pathways have been proposed in which ribo-, arabino- and threo-nucleotides would have shared common precursors (4, 5) (Figure 1). In these proposed pathways, cyanamide and glycolaldehyde react to yield 2-aminooxazole (2AO), which subsequently reacts with glyceraldehyde to form ribose aminooxazoline (RAO) and arabinose aminooxazoline (AAO), the precursors to the five-carbon sugar ribo- and arabino-pyrimidine nucleosides, respectively (4). Meanwhile, 2AO can react with a second glycolaldehyde to form threose aminooxazoline (TAO), the precursor to the four-carbon sugar threo-pyrimidine nucleosides (5). The resulting pyrimidine nucleosides can then undergo phosphorylation to generate nucleotides (6) which can then can be activated as imidazolides (7, 8). These highly reactive mononucleotides can then participate in non-templated polymerization, generating oligonucleotides that could serve as primers and templates in subsequent nonenzymatic template-directed copying reactions.

**Figure 1.**
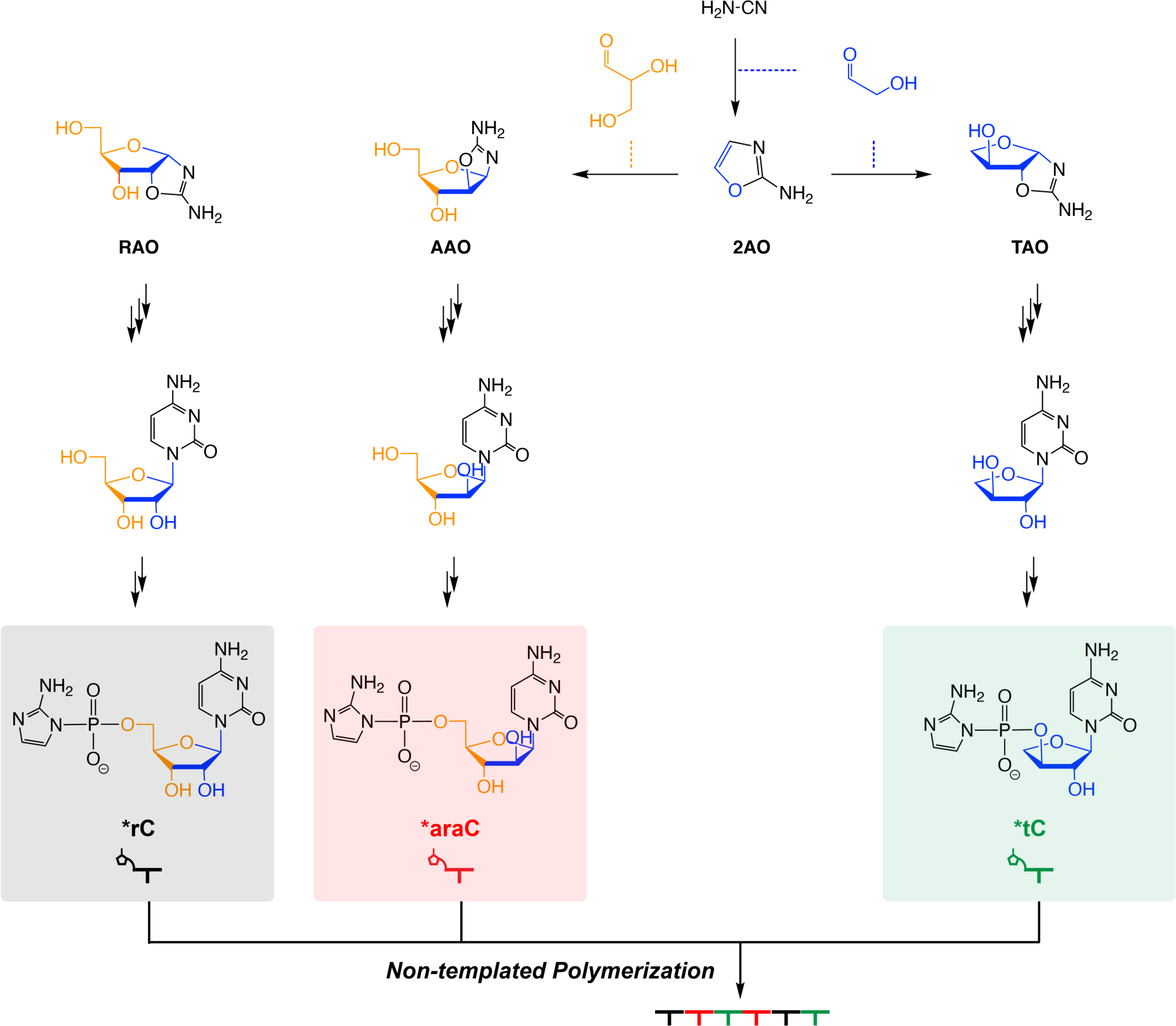
Potentially prebiotic synthetic pathway for activated ribo-, arabino-, and threo-cytidine mononucleotides. Phosphorylated and activated ribo-, arabino-, and threo-cytidine mononucleotides (*rC, *araC and *tC) can undergo non-templated polymerization to form chimeric oligonucleotides.

**Figure 2.**
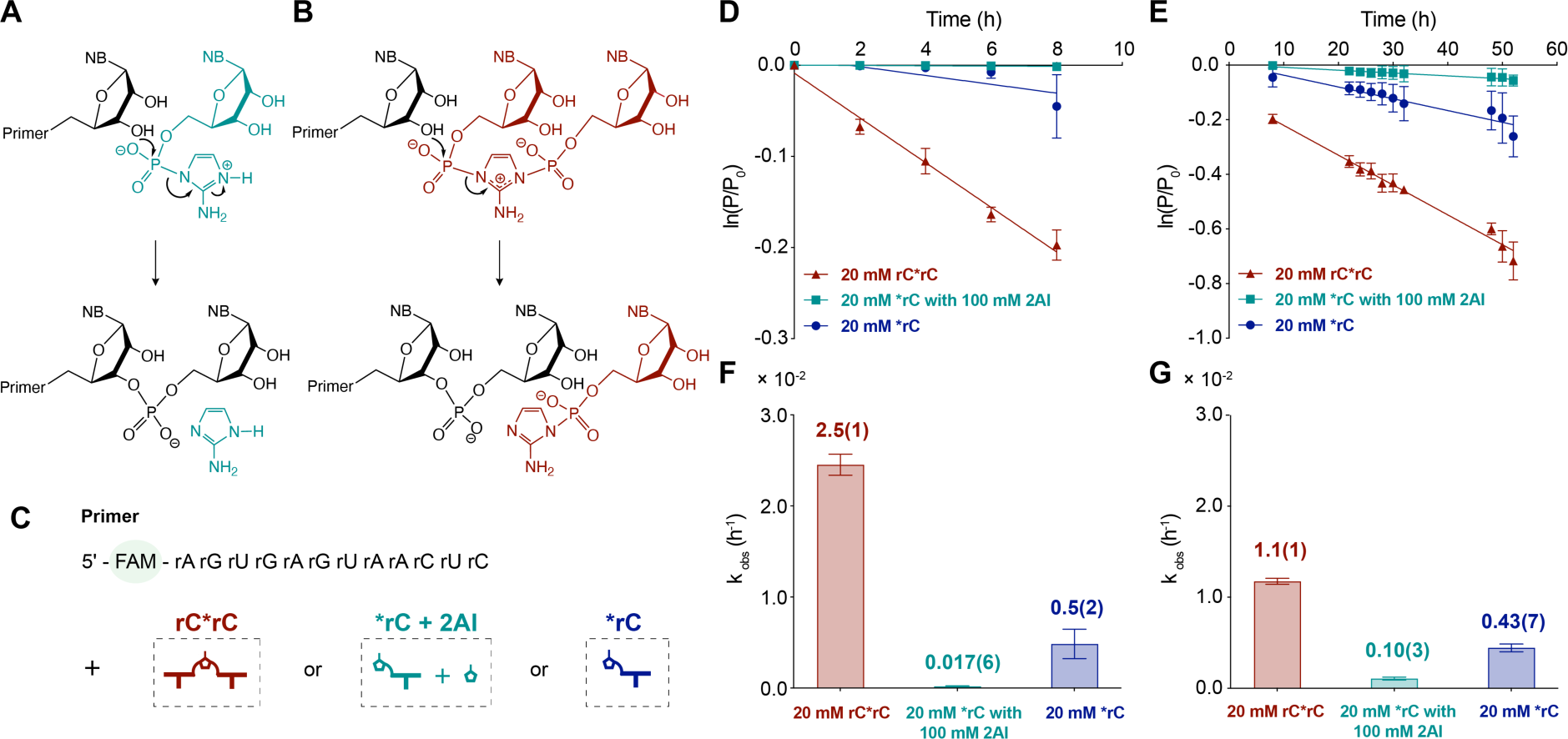
Mechanism of non-templated primer extension. Hypothesized mechanism of non-templated primer extension through (A) activated mononucleotides or (B) bridged dinucleotides. “NB”, nucleobase. Note that the scheme includes only the extension through 3ʹ-OH moieties, although nucleotides can also add to 2ʹ-OH groups. (C) Schematic representation of the non-templated primer extension using a FAM-labeled RNA primer with either bridged dinucleotides (C*C) or activated mononucleotides (*C). 2AI was added to suppress the formation of bridged dinucleotides. (D) the initial and (E) later time course. Bar plots indicate the observed pseudo-first-order reaction rates (k_obs_) calculated from (F) the initial 0-8 h and (G) the later 8-52 h time course. Reaction conditions: 1 μM FAM-labeled RNA primer (XJ-FAM-12mer, Table S1), 200 mM HEPES at pH 8.0 and 50 mM MgCl_2_ with activated species at 20 mM for rC*rC, 20 mM for *rC with 100 mM 2AI and 20 mM for *rC. Error bars represent standard deviations from the mean, n=3 replicates. Note that non-templated primer extension with 20 mM *rC and 100 mM 2AI is below the detection limit of the gel electrophoresis assay at early time points and therefore the reported initial rate might not reflect its true rate.

The non-templated polymerization of activated mononucleotides has been demonstrated on mineral surfaces such as montmorillonite (9), and also in the ice eutectic phase (10). Montmorillonite immobilizes oligonucleotides and enables successive additions of activated mononucleotides, allowing incremental elongation of the primer in the absence of a template (9). The ice eutectic phase also concentrates the solutes, enabling non-templated polymerization, while the low temperature slows down hydrolysis (10). Early studies of these processes employed imidazole-activated mononucleotides. Recent developments have shown that 2-aminoimidazole (2AI) activated nucleotides can enhance nonenzymatic template copying (11) because they spontaneously form highly reactive 5ʹ-5ʹ imidazolium bridged dinucleotides (12) that are the substrates for the predominant pathway for template-directed primer extension (13). In addition, the activating group 2AI shares a common and potentially prebiotic synthetic pathway with the precursor of nucleotide synthesis, 2AO (14).

The effect of sugar heterogeneity in the primordial pool of activated nucleotides remains a major question to be addressed en route to the RNA World. Through kinetic and crystallographic studies, template-directed copying chemistry has been shown to favor RNA synthesis (15–17). The conformation of the 3′-terminal nucleotide of the primer and the identity of the incoming bridged dinucleotides are the major determinants of the kinetics of template-directed primer extension. However, sugar heterogeneity has rarely been studied in the non-templated reactions that must have occurred to produce the first primordial templates. Is there any selectivity in non-templated reactions, given that such a reaction would be free from constraints arising from template binding? Could the structural differences between ribo-, arabino-, and threo-nucleotides influence their reactivity, shaping the composition of prebiotic oligomers? We have attempted to gain insight into the likely distribution of oligonucleotide products by simulating these conditions through competition experiments with mixtures of nucleotide substrates.

In this study, we first examined the mechanism of non-templated primer extension and identified imidazolium-bridged dinucleotides as the most reactive substrates. Subsequent kinetic analyses using 2AI-activated ribo-, arabino-, and threo-nucleotides showed that ribonucleotides and arabino-nucleotides were incorporated into a growing primer at similar rates, while threo-nucleotides displayed lower reactivity. We then combined these nucleotides to mirror the heterogenous prebiotic environment, and used LC-MS to investigate the products of non-templated primer extension. Again, arabino-nucleotides and ribonucleotides were incorporated to similar extents, whereas threo-nucleotides were outcompeted. Our findings demonstrate an inherent bias against the incorporation of threo-nucleotides during non-templated polymerization.

## MATERIAL AND METHODS

### General Information

#### Materials

Reagents and solvents were obtained from Fischer Scientific, Sigma-Aldrich, Alfa Aesar, Acros Organics and were used without further purification unless otherwise noted. RNA and DNA oligonucleotides were purchased from Integrated DNA Technologies or synthesized in-house using the Expedite 8909 DNA/RNA synthesizer. Further information on oligonucleotide synthesis is included in the supplementary data.

#### Nuclear Magnetic Resonance (NMR)

^1^H and ^31^P NMR spectra were acquired on a Varian Oxford AS-400 NMR spectrometer (400 MHz for ^1^H, 162 MHz for ^31^P) at 25 °C. ^1^H NMR spectra were referenced using sodium 2,2-dimethyl-2-silapentane-5-sulfonate (DSS) as internal standard (0 ppm at 25 °C).

#### Low-resolution Mass Spectrometry (LRMS)

All samples were diluted to 200 μM in 50% (v/v) acetonitrile immediately prior to analysis. The spectra were obtained by direct injection on an Esquire 6000 mass spectrometer (Bruker Daltonics), which was operated in the alternating ion mode.

#### High-resolution Liquid Chromatography Mass Spectrometry (HR LC-MS)

The eluted samples were separated and analyzed using an Agilent 1200 High-performance Liquid Chromatography (HPLC) coupled to an Agilent 6230 Time-of-flight Mass Spectrometry (TOF MS) equipped with a diode array detector. The samples were separated by IP-RP-HPLC on a 100 mm × 1 mm (length × i.d.) Xbridge C18 column with a 3.5 μm particle size (Waters, Milford, MA). The samples were eluted between 2.5 and 15% methanol in 200 mM 1,1,1,3,3,3-hexafluoro-2-propanol with 1.25 mM triethylamine at pH 7.0 over 16 minutes with a flow rate of 0.1 mL/min at 50 °C. The samples were analyzed in negative mode from 239 m/z to 3200 m/z with a scan rate of 1 spectrum/s.

### Synthesis and Characterization of 2-aminoimidazole Activated Mononucleotides and 5ʹ-5ʹ Imidazolium-bridged Dinucleotides

#### Arabinose guanosine nucleoside (araG) phosphorylation

The Yoshikawa method was used to phosphorylate the 5ʹ-hydroxyl of araG nucleoside (18). To a pre-chilled mixture of 9-(β-D-arabinofuranosyl)guanine (araG, 1 equiv.) in trimethyl phosphate (OP(OMe)_3_, 0.1 M with respect to the nucleoside) was added phosphoryl chloride (POCl_3_, 4 equiv.) and a trace amount of H_2_O (7.5 μL) under vigorous stirring. The resultant mixture was allowed to stir at 0 °C. After complete solubilization of the nucleoside, four portions of *N,N*-diisopropylethylamine (DIPEA, 0.5 equiv. each) were added dropwise at 20 min intervals. The reaction was monitored by LRMS. Once the starting material disappeared, the reaction was quenched using an aqueous solution of 1 M triethylamine-bicarbonate (TEAB, 5 volumes, pH 7.5). Any precipitate observed after the quench was re-solubilized using a minimal amount of acetonitrile. The products were purified by reverse phase flash chromatography with a 50 g C18Aq column. The desired product was separated from other compounds over 10 column volumes (CVs) of 0–15% acetonitrile in 2 mM aqueous TEAB buffer (pH 7.5) with a flow rate of 40 mL/min. Fractions containing the product were collected and lyophilized under high vacuum level at room temperature. The 9-(β-D-arabinofuranosyl)guanine monophosphate was used for activation without further purification.

#### Threose cytidine nucleoside (tC) phosphorylation and deprotection

The benzoyl protecting groups on the 2ʹ-hydroxyl and amine enabled selective phosphorylation of the 3ʹ-hydroxyl. The reaction is outlined in Scheme S1. To azeotropically dried (toluene 3x, 2.5 mL each) tC (1 equiv.) and bis(2-cyanoethyl)-N,N-disopropylphosphoramidite in acetonitrile (0.1 mL, 1.25 equiv.) was added a solution of 5-(ethylthio)-1H-tetrazole (ETT, 2.5 equiv.) in acetonitrile dropwise. The mixture was stirred for 1 h. The reaction was monitored by thin layer chromatography (TLC). Meta-chloroperoxybenzoic acid (mCPBA, 3.0 equiv.) was added and the mixture was stirred for another 5 mins, monitored by TLC. The reaction mixture was diluted with 15 mL DCM, transferred to a separatory funnel and 10 mL aqueous sodium bicarbonate (NaHCO_3_) was added. The aqueous layer was extracted with dichloromethane (DCM) two times and the combined organic extracts were washed with brine. The extracts were dried with sodium sulfate and concentrated by rotary evaporator. The compound was dissolved in 0.5 mL water. Following the addition of 20 mg mCPBA, the reaction was well mixed. 8 mL of ammonium hydroxide (NH_4_OH) was then added. The progress of the deprotection reaction was monitored by LRMS. A heat gun was used to evaporate the ammonia before concentrating the product in a rotary evaporator. The dried compound was dissolved in 5 mL 200 mM TEAB at pH 9 and purified by reverse phase flash chromatography with a 50 g C18Aq column. The desired product was separated from other compounds over 10 CVs of 0–5% acetonitrile in 2 mM TEAB buffer (pH 7.5) with a flow rate of 40 mL/min. The collected fractions were measured by LR-MS and lyophilized for further activation.

#### Synthesis and characterization of 2-aminoimidazole activated mononucleotides (*N)

The activation of NMPs followed a previously reported procedure (11). The activation of threose cytidine and guanosine monophosphate (tCMP and tGMP) was similar to the NMP activation detailed above except for the following steps: 1) tCMP or tGMP (1.0 equiv.), 2AI·HCl (10 equiv.) and TPP (0.5 equiv.) were suspended in 30 mL DMSO under Ar, and 2) The collected fractions were pH adjusted to 9 by 1M NaOH before lyophilization. The detailed characterizations (NMR and HR-MS) are included in the supplementary data.

#### Synthesis and characterization of 5ʹ-5ʹ 2-aminoimidazolium-bridged dinucleotides (N*N)

The synthesis of N*N followed a previously reported protocol (19). The detailed characterizations (NMR and HR-MS) are included in the supplementary data.

### Non-templated Primer Extension Reactions in the Aqueous Phase

#### Non-templated primer extension

Non-templated primer extension reactions were performed at 1 μM primer, 20 mM activated mononucleotides, 200 mM HEPES at pH 8.0 and 50 mM MgCl_2_. At each time point, 1 μL of the reaction sample was added to 19 μL of quenching buffer containing 7M urea, 1x TBE, 100 mM EDTA.

#### Urea-PAGE analysis

A fluorophore-labeled primer was utilized to visualize primer extension by polyacrylamide gel electrophoresis (PAGE). Primers were either 12- or 6-nucleotides long. Primer extension products were resolved by 20% PAGE with 7 M urea, in 1x TBE gel running buffer. For extension reactions using a 6-mer primer, the reactions were desalted by ion pairing reverse phase (IP-RP) purification using C18 ZipTip pipette tips (Millipore, Billerica, MA) to avoid smearing of bands in PAGE. The tips were wetted with 100% (v/v) acetonitrile and equilibrated with 100 mM TEAA prior to sample binding. Extensive washing with 100 mM TEAA, then with LC-MS grade water, was followed by elution in 50% (v/v) acetonitrile. The eluates were dried in a centrifugal vacuum concentrator to remove acetonitrile, followed by resuspension in 10 μL of 7 M urea and 1x TBE for sample loading. An increased acrylamide concentration (22%) was used for improved separation of smaller molecules.

The gels were scanned with an Amersham Typhoon RGB Biomolecular Imager (GE Healthcare Life Sciences) to visualize the fluorophore-labeled primer and extended primer bands. The relative band intensities were quantified using ImageQuant TL software. The rate of the extension was determined from the linear least-square fits of the data.

### Spontaneous Air-Drying Experiments, Competition Experiments and Subsequent LC-MS Analysis

#### Spontaneous air-drying experiments

PCR tubes containing 10 μL of non-templated primer extension reactions were prepared with 20 mM activated mononucleotides, 100 μM 6-mer primer, 200 mM HEPES at pH 8.0, and 50 mM MgCl_2_. The reactions were allowed to dry spontaneously under ambient air, resulting in clear pastes (Figure S1). The reaction pastes were then quenched at 24 h by adding 58 μL of quench buffer containing 100 mM EDTA at pH 8.0. Spontaneous air-drying significantly accelerated the non-templated primer extension, thereby generating enough primer +1 products for the following LC-MS experiments.

#### Spontaneous air-drying experiments in the presence of complementary oligomers

The primer/complementary-oligomer duplexes were prepared in an annealing buffer at 2.2 times the final concentration. Solutions containing 217 μM primer (XJ-8, Table S1), 1630 μM complementary oligomers (XJ-6mer-rc, Table S1), 22 mM HEPES at pH 8.0, 22 mM NaCl, and 0.44 mM EDTA at pH 8.0 were heated at 95 °C for 30s and then slowly cooled to 25 °C at a rate of 0.1 °C/s in a thermal cycler machine. The annealed products were then diluted into the primer extension reaction buffer to give final concentrations of 100 μM primer, 750 μM template, 200 mM HEPES at pH 8.0, and 50 mM MgCl_2_.

#### Competition experiments

In competition experiments, unlabeled primer was mixed with activated ribo-, arabino- and threo-nucleotides at the indicated ratio and left to air-dry spontaneously (reaction conditions as outlined in the spontaneous air-drying experiments). For an equimolar mixture (*rC:*araC:*tC = 1:1:1), the total of 20 mM activated mononucleotides consisted of 6.7 mM each of *rC, *araC and *tC. For the *rC:*araC:*tC = 10:1:1 ratio, the total of 20 mM activated mononucleotides consisted of 16.7 mM *rC, 1.7 mM *araC and 1.7 mM *tC. For the experiment shown in Figure 4, the primer was XJ-5 (Table S1) and for Figure 6 was XJ-10 (Table S1).

#### Primer design for competition experiments

To improve the product separation in LC-MS, we used a 6-nt long 5ʹ-OH primer instead of a fluorophore-labeled primer. In Ion Pairing Liquid Chromatography (IP-LC), separation is achieved through the difference in electrostatic affinities of charged molecules for a charged stationary phase (20). However, separating RNA oligomers of the same length can be challenging. To separate the +1 products ending in a ribo-, arabino-, or threo-nucleotide, we used shorter RNA oligomers with lower total charge. Furthermore, replacing a fluorophore-labeled primer with an unlabeled primer decreased the total charge and improved the separation of same-length RNA oligomers with different terminal nucleotides.

#### LC-MS sample preparation

Reaction mixtures to be subjected to LC-MS analysis were quenched with 100 mM EDTA at pH 8.0. The quenched reactions (60 μL) were aliquoted into 10 uL fractions for desalting by ion pairing reverse phase (IP-RP) purification on C18 ZipTip pipette tips (Millipore, Billerica, MA) as previously described. The eluates were combined into an HPLC vial insert and dried in a centrifugal vacuum concentrator. Samples were resuspended in LC-MS grade water prior to injection in HR LC-MS.

#### LC-MS analysis

Extracted compound chromatograms were generated using the Agilent MassHunter Qualitative Analysis software (B.07.00). For competition experiments, generated compound lists were matched with the calculated masses of all possible +1 non-templated primer extension products and their salt adducts. The observed and calculated masses of the relevant products are provided in Table S2 and Table S3. For the purposes of this analysis, we assumed that the same-length oligomers ending in different terminal sugars have equivalent ionization efficiencies.

## RESULTS

### Mechanism of non-templated primer extension

We hypothesized two distinct pathways for non-templated primer extension: one through the direct reaction of activated mononucleotides (Figure 2A) and the other through imidazolium bridged dinucleotides (Figure 2B). The activated mononucleotide pathway proceeds when the terminal hydroxyl group of the primer (3ʹ-OH or 2ʹ-OH) attacks the phosphorus atom of an activated mononucleotide, displacing 2-aminoimidazole (2AI) as the leaving group (Figure 2A). In contrast, the bridged dinucleotide pathway is characterized by the hydroxyl group attacking the phosphorus atom of an imidazolium bridged dinucleotide, displacing an activated mononucleotide as the leaving group (Figure 2B). These highly reactive dinucleotides form spontaneously by the reaction of two activated mononucleotides (21, 22). While bridged dinucleotides are known to drive templated nonenzymatic copying (13), their role in non-templated primer extension remains unclear due to the lack of stabilizing base pair interactions with the template.

To elucidate the mechanism of non-templated primer extension, we conducted comparative kinetic analyses of primer extension reactions using activated cytidine mononucleotides (*rC) or 5ʹ-5ʹ imidazolium bridged dinucleotides (rC*rC) (Figure 2C). The reaction with 20 mM pre-formed purified bridged rC*rC started off fast (Figure 2D), but then slowed down (Figure 2E), presumably due to hydrolysis and an approach to a steady state equilibrium mixture of *rC and rC*rC. In contrast, the reaction with 20 mM *rC started off very slowly, but speeded up with time, again presumably due to the formation of an equilibrium mixture of *rC and rC*rC. Because *rC can spontaneously react to form bridged dinucleotides in situ, we added excess 2AI to reactions containing *rC to minimize the accumulation of bridged dinucleotides (12), which resulted in a sustained slow rate of primer extension (Figure 2D-G). To obtain a valid comparison of reaction rates with *rC and rC*rC, we first measured initial rates with 20 mM rC*rC (k = 2.5(1)× 10^-2^ h^-1^) and with 20 mM *rC (k = 5(2) × 10^-3^ h^-1^). To eliminate the effect of rC*rC formation in the reaction with *rC, we measured the reaction rate with 20 mM *rC in the presence of excess 2AI, and observed an even lower rate of extension (< 0.2 × 10^-3^ h^-1^) (Figure 2F). We observed a similar pattern of reactivity with activated guanosine nucleotides (Figure S2). The primary role of bridged dinucleotides in non-templated primer extension is evident from our observation that the initial reaction rates are substantially elevated in reaction mixtures containing bridged dinucleotides (rC*rC and *rC alone) compared to those with minimal bridged dinucleotide formation (*rC + 2AI) (Figure 2D, F). The reactivity difference stems from the fact that bridged dinucleotides do not require protonation of the leaving group (Figure 2B), whereas activated mononucleotides require protonation (13) (Figure 2A). This is consistent with a comparison of their hydrolysis rates: the rate of hydrolysis of rC*rC is 0.105(7) h^-1^ while the hydrolysis of *rC is significantly lower at 1.96(6) × 10^-3^ h^-1^ at pH 8.0 (Figure S3). We conducted non-templated primer extension experiments as a function of pH to determine if increased protonation of the leaving group at lower pH would accelerate the reaction rate. Our results show that the rates for the reactions involving 5 mM *rG and 25 mM 2AI at pH 7, 8, and 9 are 1.9(3) × 10^-4^, 5.3(6) × 10^-4^, and 8.1(4) × 10^-4^ h^-1^, respectively (Figure S4). The decreased reaction rate at lower pH is likely due to decrease in deprotonation of the 3ʹ-OH of the primer.

We followed the distribution of activated monomers and bridged dinucleotides as a function of time under primer extension reaction conditions by ^31^P NMR (Figure S5). In a reaction mixture that initially contained 20 mM bridged dinucleotides, the concentration of bridged dinucleotides declined due to hydrolysis, with a half-life of 7 h (Figure S3A). In contrast, a reaction initiated with 20 mM of *rC alone showed a rapid initial increase in the concentration of rC*rC to an expected equilibrium value of approximately 3 mM, followed by a long slow decline to 2.5 mM by 8 h. In contrast, the mixture of 20 mM *rC with 100 mM 2AI contained less than 0.25 mM of bridged dinucleotides at all times. In all cases the concentration of activated mononucleotides decreased very slowly with time, with an estimated half life of ∼350 h (Figure S3B). This concentration of bridged dinucleotides in these different reaction mixtures correlates well with the initial rates of primer extension in reactions initiated with either bridged dinucleotides or activated mononucleotides. After the initial equilibration period, the hydrolysis of bridged dinucleotides into activated mononucleotides and unactivated nucleotide monophosphates results in a slowly declining reaction rate (Figure 2E,G). Consequently, kinetic and NMR data support the conclusion that bridged dinucleotides are the primary substrates for non-templated primer extension.

### Comparison of non-templated primer extension with activated ribo-, arabino- and threo-nucleotides

Given that ribo-, arabino- and threo-nucleotides might all be synthesized together in a prebiotic environment, we wanted to evaluate whether the intrinsic chemical and structural differences between these nucleotide analogs would affect their incorporation into oligonucleotides. To achieve this, we added activated ribo-, arabino- and threo-cytidine mononucleotides (*rC, *araC, and *tC) separately to the primer (Figure 3A). We used activated mononucleotides because they naturally form bridged dinucleotides and their rate of reaction is only modestly lower than pure bridged dinucleotides in an extended time course (k = 4.4(4) × 10^-3^ h^-1^ versus 1.2(1) × 10^-2^ h^-1^). The non-templated primer extension rate using *rC was found to be slightly faster at 1.9(1) × 10^-3^ h^-1^ than the extension rate observed when only *araC was used, which was 1.5(1) × 10^-3^ h^-1^ (Figure 3B, C). However, *tC did not yield detectable extension. A similar trend was observed with activated ribo-, arabino-, and threo-guanosine mononucleotides (*rG, *araG, and *tG) (Figure 3D-F). The *rG resulted in the fastest extension rate of 2.2(1) × 10^-3^ h^-1^, closely followed by *araG at 1.8(1) × 10^-3^ h^-1^. *tG had the slowest extension rate at 3.6(8) × 10^-4^ h^-1^, which was approximately 6-fold slower than that with *rG (Figure 3F).

**Figure 3.**
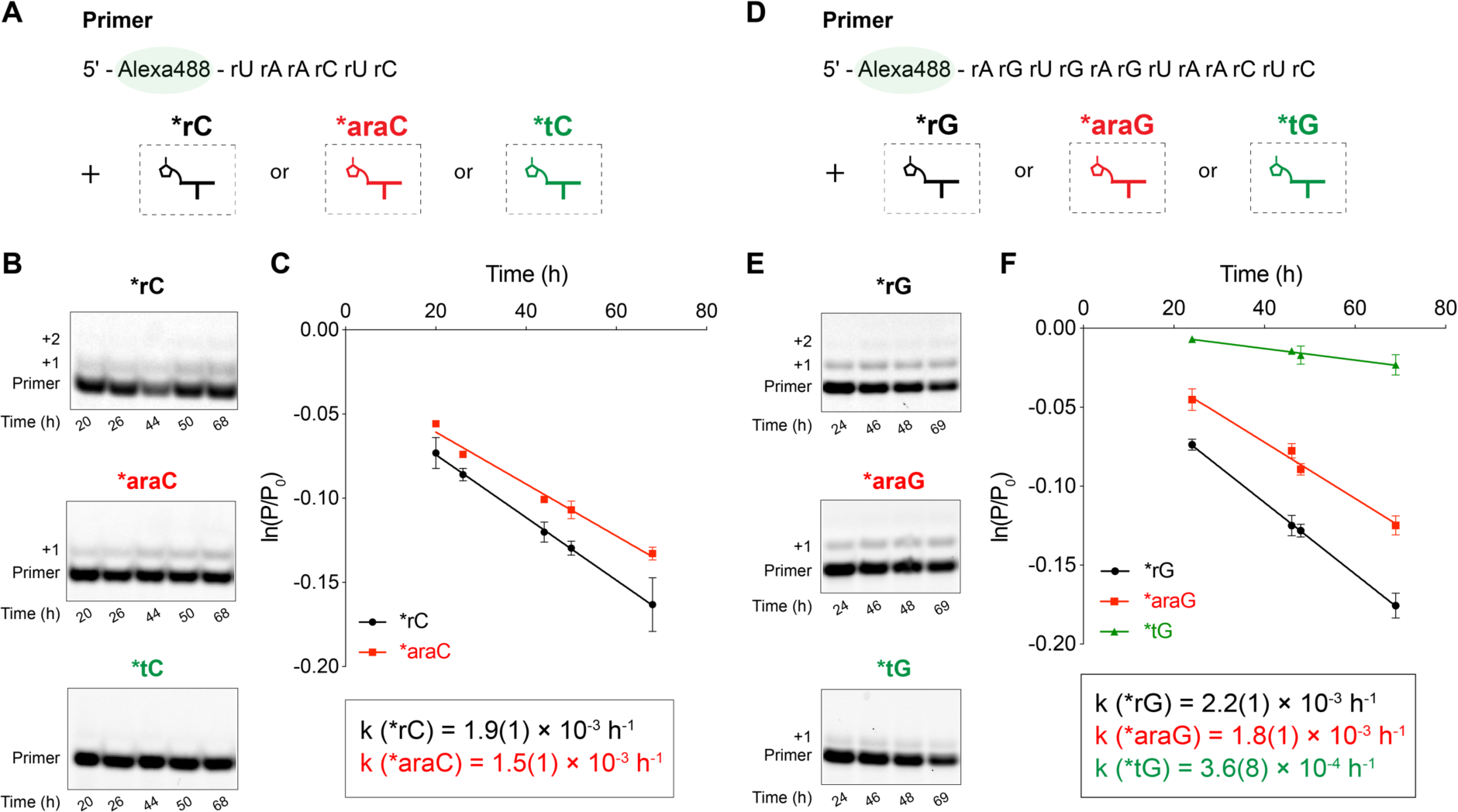
The intrinsic reactivities of activated ribo-, arabino- and threo-nucleotides in non-templated primer extension with (A-C) cytidine and (D-F) guanosine mononucleotides. (A, D) Schematic representation of non-templated primer extension reaction using an Alexa488-labeled RNA primer with the addition of activated ribo-, arabino-, or threo-nucleotides, respectively. (B, E) Gel electrophoresis images. (C, F) Kinetic analysis with observed pseudo-first-order reaction rates (k_obs_). Reaction conditions: 1 μM primer (XJ-Alexa-6mer or XJ-Alexa-12mer, Table S1), 200 mM HEPES at pH 8.0 and 50 mM MgCl_2_ with 20 mM of *rC or *rG, *araC or *araG, and *tC or *tG, respectively. Error bars represent standard deviations from the mean, n=2 replicates.

**Figure 4.**
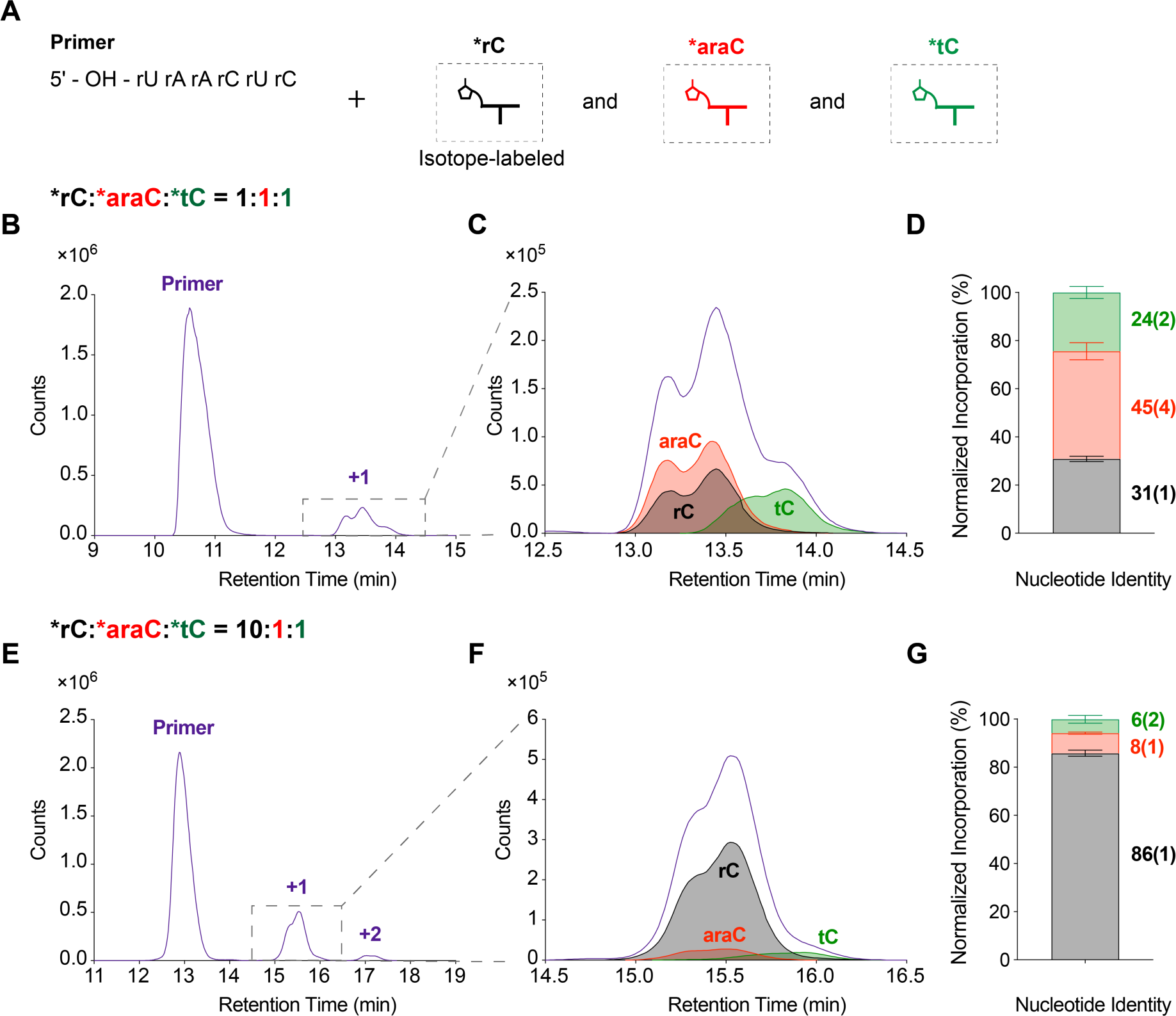
Incorporation of ribo-, arabino-, and threo-nucleotides in competition experiments. (A) Schematic representation. (B-D) *rC:*araC:*tC = 1:1:1 and (E-G) 10:1:1 competition experiments with (B, E) total compound chromatogram (TCC) of the +1 extended products (purple); (C, F) overlay of extracted compound chromatograms (ECC) of the +1 extended products ending in rC (black), araC (red) and tC (green); (D, G) stacked barplot representing the normalized incorporation of different nucleotides. Error bars represent standard deviations from the mean, n=6 replicates.

Arabino-nucleotides are structurally similar to ribonucleotides, with the only difference being the stereochemistry of the sugar 2’-OH group. Kim *et al.* (15) showed that this difference does not impact the stability or formation of bridged dinucleotides under primer extension conditions. The rates of hydrolysis of bridged ribonucleotides (rA*rA) and arabino-nucleotides (araA*araA) are similar, at k = 0.31 h^-1^ and k = 0.33 h^-1^, respectively. On the other hand, threo-nucleotides lack the 5’-methylene carbon found in ribo-nucleotides, resulting in a backbone repeat unit that is one atom shorter in TNA than in RNA. In addition, the rate of formation of bridged threo-nucleotides (tC*tC) is 8.8 × 10^-4^ h^-1^ mM^-1^ (17), while that of bridged ribonucleotides (rC*rC) is 4.5 × 10^-3^ h^-1^ mM^-1^ (12). The formation of 3ʹ-3ʹ imidazolium-bridged threo-nucleotides is impeded by steric hindrance, in contrast to the facile formation of 5ʹ-5ʹ imidazolium-bridged ribonucleotides (Figure S6). This increased steric hindrance accounts for the observed five-fold reduction in the rate of formation of bridged threo-nucleotides. In the non-templated primer extension reaction, the slower rate of formation bridged threo-nucleotides and the resulting lower steady state level of bridged dinucleotides would be expected to result in a slower primer extension rate as non-templated primer extension primarily occurs via bridged dinucleotides.

### Non-templated primer extension with mixture of activated ribo-, arabino- and threo-nucleotides

To simulate the diverse chemical pool of activated nucleotides that could have co-existed on the early Earth, we proceeded to study the products of non-templated primer extension using a mixture of activated ribo-, arabino- and threo-nucleotides. To analyze the product distribution from these competition reactions, we used liquid chromatography-mass spectrometry (LC-MS). To distinguish the otherwise isobaric ribo-nucleotides and arabino-nucleotides, we used stable-isotope labeled (13C, 15N) ribonucleotides.

To accelerate the intrinsically slow non-templated reactions, we used a spontaneous air-drying method which reduces the time required to observe appreciable product yield. Spontaneous air-drying is a prebiotically plausible scenario driven by environmental factors such as fluctuations in temperature and humidity, precipitation and evaporation, day-night cycles and the presence of hydrothermal vents (23). Dry-down has been extensively used to model potentially prebiotic conditions in the field of prebiotic chemistry (24, 25). We left the reaction mixture in an uncapped tube on the bench top to allow water to evaporate for 24 hours (Figure S1). We observed minimal variability within the same set of experiments by this method (Figure S7). However, it is important to note that variability across different sets of experiments could be significant due to the variable flow rate and moisture level of the ambient air (Table S4). The heterogeneity of the early Earth’s environment would likely provide more variable conditions than found in a confined system or an aqueous environment (24). To maximize the amount of +1 products available for accurate LC-MS characterization, we screened a range of oligonucleotide concentrations for air-drying experiments and chose 100 μM as the starting concentration for spontaneous air-drying and competition experiments (Figure S8).

#### Incorporation of ribo-, arabino-, and threo-nucleotides in competition experiments

We examined non-templated primer extension in the presence of different ratios of ribo-, arabino-, and threo-nucleotides and analyzed the products of primer extension by LC-MS (Figure 4A). In a mixture of activated mononucleotides in a 1:1:1 ratio (*rC:*araC:*tC = 1:1:1), we observed both the primer and the +1 products in the total compound chromatogram (TCC) (Figure 4B). To distinguish between the +1 extended products for each mononucleotide, we used extracted compound chromatograms (ECC) for the expected extended products and overlaid them with the +1 product region of the TCC (Figure 4C). We confirmed that each peak in the ECC corresponded to the expected +1 products (Table S2). We observed the expected mass-to-charge ratios (m/z) of the different +1 products and their salt adducts (Figure S9), demonstrating that each species could be easily distinguished. We calculated the percentage of each nucleotide in the observed +1 products (Figure 4D, Table S4A), revealing that arabino-nucleotide incorporation (44.7 ± 3.5 %) was the highest, followed by ribonucleotide (30.9 ± 1.1 %) and threo-nucleotide incorporation (24.4 ± 2.5 %).

We performed an additional competition experiment with *rC, *araC, and *tC at a ratio of 10:1:1 to simulate the effect of enrichment for ribonucleotides during prebiotic synthesis (Figure 1). It has been observed that ribose aminooxazoline (RAO), the precursor to ribonucleotides, can crystallize out, leaving other aminooxazolines in solution (26). Nucleotides derived from crystallized RAO would be predominantly ribonucleotides. Furthermore, homochiral RAO crystals can form through crystallization on magnetite (Fe_3_O_4_) surfaces, which induce chiral symmetry-breaking (27). The crystallization of enantiopure RAO, in turn, triggers the avalanche magnetization of magnetite, forming a closed feedback loop between chiral molecules and magnetite surfaces (28). Alternatively, enantiopure RAO can be obtained through physical and chemical amplification processes using enantioenriched amino acids (29). Taken together, these phenomena suggest that prebiotic mixtures of nucleotides might have been strongly enriched in ribonucleotides. We therefore sought to investigate how changes in the stoichiometry of prebiotic mixtures, particularly the enrichment of ribonucleotides, would influence the composition of the products in downstream non-templated polymerization.

In competition reactions initiated with *rC:*araC:*tC at a 10:1:1 ratio, we observed primer, +1 and +2 products (Figure 4E). By overlaying the TCC of the +1 products with the ECC of the expected products (Figure 4F), we found that ribonucleotide incorporation was the highest (85.8 ± 1.3 %), followed by arabino-nucleotide (8.4 ± 0.4 %), and then threo-nucleotide incorporation (5.8 ± 1.6 %) (Figure 4G, Table S4A). The ratio of incorporated ribonucleotide to arabino-nucleotide was ∼10 to 1 in the +1 products, while threo-nucleotide incorporation was slightly disfavored. This result suggests that the composition of non-templated extension products will approximately mirror the input ratios of activated ribo-, arabino- and threo-mononucleotides, with a moderate selection bias against threo-nucleotides. Consequently, the enrichment of RAO in prebiotic mixtures would be manifested in the composition of the products of subsequent non-templated reactions. These consecutive steps, occurring during prebiotic synthesis and non-templated reactions, could therefore synergistically exert a selection pressure favoring ribonucleotide incorporation.

#### Non-templated reaction at both internal and terminal hydroxyls

The chromatograms shown in Figure 4 exhibited a double peak pattern for +1 products in both the TCC and the ECC. To trace the origins of the two peaks in the chromatograms, we considered all possible reaction sites in an RNA oligomer. Nucleotides contain two major nucleophilic groups, of which the 2ʹ-hydroxyl group is more reactive than the 3ʹ-hydroxyl (30, 31). During non-templated primer extension with activated species, oligonucleotides with mixed 2ʹ-5ʹ and 3ʹ-5ʹ phosphodiester linkages can form. We first examined the possibility of the double peak pattern arising from a mixture of terminal 3ʹ-5ʹ and 2ʹ-5ʹ phosphodiester linkages by co-injecting oligonucleotides synthesized with terminal 3ʹ-5ʹ and 2ʹ-5ʹ linkages from the corresponding phosphoramidites at 1:1 and 1:10 ratios. These terminal linkage regioisomers could not be resolved by LC-MS (Figure S10A, B), showing that terminal regioisomers were not responsible for the observed double peak pattern.

To narrow down the search for the cause of the double peak pattern, we performed a series of non-templated reactions with various primer constructs. We first examined the possibility of reaction at the 5ʹ-OH group of the primer by carrying out a dry-down reaction with a 5ʹ-hexynyl DNA primer and a 5ʹ-OH DNA primer, each with a terminal dideoxy nucleotide. In both cases, no extension products were observed by LC-MS (Figure S10C, D). This result suggested that at least one of the two HPLC peaks might have derived from reaction at internal 2ʹ-hydroxyls.

Further tests with modified RNA primers confirmed that the internal 2ʹ-OHs in RNA oligomers can react with activated nucleotides (Figure 5A-D, Figure S11), causing the double peak pattern. Comparing the TCC and ECC of the non-templated products of a 5ʹ-hexynyl RNA primer ending with dideoxycytosine (Figure S11B) and an all RNA 5ʹ-hexynyl primer (Figure S11C) shows that both internal and terminal hydroxyls can react under spontaneous air-drying conditions. The experimentally observed ratio of internal reaction to terminal extension was ∼ 1:1.6 (Figure 5D). Since there are five internal hydroxyls, but only two terminal hydroxyls (Figure S11A), on average the internal hydroxyls must have only ∼25% of the reactivity of the terminal hydroxyls (Table S4B, Figure 5D). In the presence of complementary short oligomers, total internal hydroxyl reactivity was reduced to 21(1)% (Figure 5E-H), corresponding to an average ratio of internal to terminal hydroxyl reactivity of 10(1)% (Table S4B, Figure 5H). This observation suggests that internal 2ʹ-OH modification would be reduced in the presence of oligonucleotides derived from both strands of a circular sequence, as hypothesized in the Virtual Circular Genome Model for primordial RNA replication (32).

**Figure 5.**
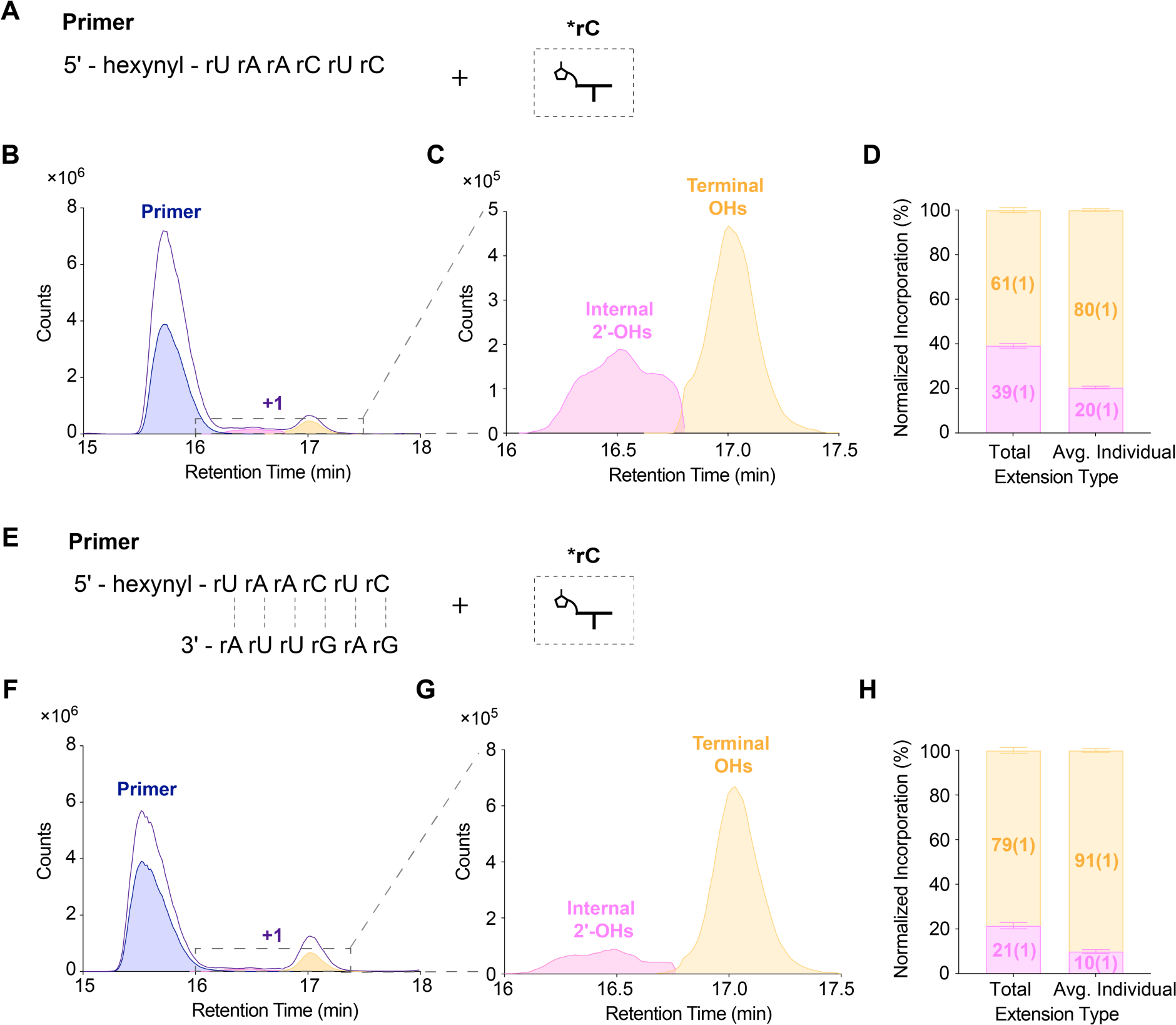
Complementary oligomers reduce the reactivity of internal 2ʹ-OHs. Non-templated primer extension of a 5ʹ-hexynyl RNA primer in the absence (A-D) or presence (E-H) of a complementary oligomer. (A, E) Schematic representation; (B, F) overlay of TCC (purple) and ECC (blue: primer; pink: internal 2ʹ-OHs; yellow: terminal OHs); (C, G) overlay of ECC corresponding to reaction of internal 2ʹ-OHs or terminal OHs; (D, H) stacked barplots representing the normalized incorporation at internal 2ʹ-OHs or terminal-OHs in total vs. average individual reaction. Error bars represent standard deviations from the mean, n=6 replicates.

**Figure 6.**
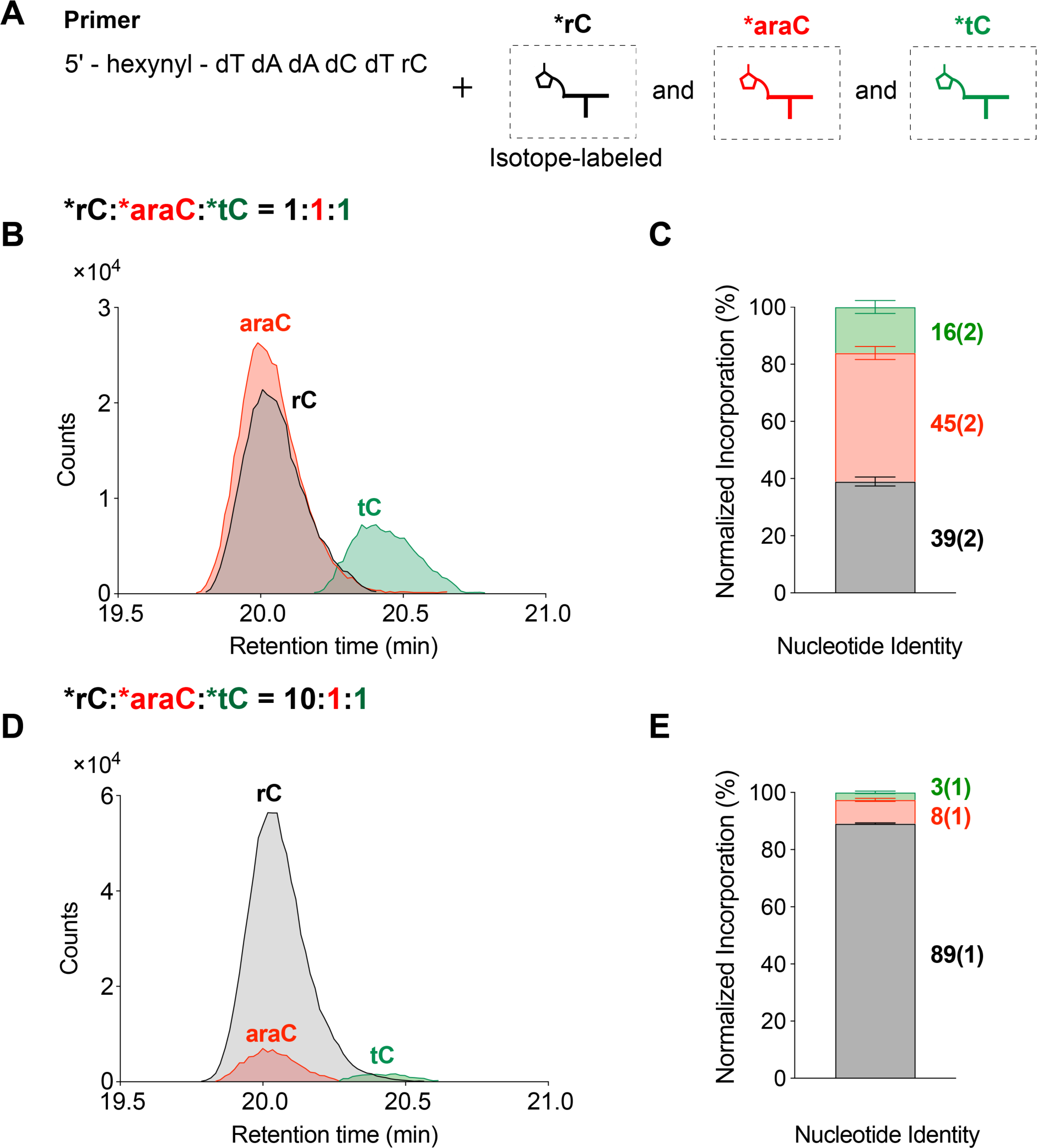
Competition experiments with a 5ʹ-hexynyl DNA primer with a 3ʹ-terminal ribonucleotide. (A) Schematic representation. Competition experiments with (B-C) *rC:*araC:*tC = 1:1:1 and (D-E) 10:1:1: (B, D) overlay of the ECC of the +1 extended products ending in rC (black), araC (red), and tC (green); (C, E) stacked barplot representing the normalized incorporation percentage of different nucleotides. Error bars represent standard deviations from the mean, n=6 replicates.

#### Competition experiments with a 5ʹ-hexynyl DNA primer with a 3ʹ-terminal ribonucleotide

To investigate the non-templated addition of ribo-, arabino- and threo-nucleotides specifically to the primer terminus, we carried out dry-down reactions using a 5ʹ-hexynyl DNA primer with a 3ʹ-terminal ribonucleotide, so that only the terminal diol is available for reaction (Figure 6A). We then analyzed the observed masses of +1 products (Table S3). The m/z profiles of all +1 products and their salt adducts were well separated, allowing us to distinguish the different species (Figure S12). ECCs generated from these LC-MS datasets showed no peak doublet pattern in any of the three +1 products corresponding to ribo-, arabino-, and threo-nucleotide incorporation, at input ratios of *rC:*araC:*tC of 1:1:1 (Figure 6B, C) or 10:1:1 (Figure 6D, E). Consistent with the trends observed in the previous competition experiments, at a 1:1:1 input ratio, the modified primer results also showed that ribonucleotides and arabino-nucleotides exhibited comparable levels of incorporation (39.0 ± 1.6 % and 45.0 ± 2.3 %, respectively). As before, threo-nucleotide incorporation was significantly reduced (16.1 ± 2.3 %) (Figure 6C, Table S4C). To simulate the enrichment of ribonucleotides in prebiotic mixtures, we again changed the input ratio of *rC:*araC:*tC to 10:1:1. Upon the analysis of the resulting +1 products, ribonucleotide incorporation was predominant (89.2 ± 0.2 %), followed by arabino-nucleotide (8.3 ± 0.5%), and then by threo-nucleotide incorporation (2.6 ± 0.4%) (Figure 6E, Table S4C).

## DISCUSSION

Potentially prebiotic synthetic routes suggest that ribo-, arabino-, and threo-nucleotides may have been synthesized together on the early Earth. Once activated these nucleotides could have co-polymerized to form oligomers, resulting in a heterogenous mixture of strands capable of taking part in template-directed nonenzymatic copying reactions and potentially cycles of replication (Figure 7). Despite the significance of non-templated polymerization in understanding the emergence of RNA from prebiotic mixtures, this step has been poorly studied. We have investigated the mechanism of non-templated primer extension and its behavior with activated ribo-, arabino- and threo-nucleotides. In addition, we performed one-pot competition experiments with mixtures of these activated nucleotides at varying input ratios to simulate the content of prebiotic mixtures and to monitor the composition of the resulting non-templated primer extension products.

**Figure 7.**
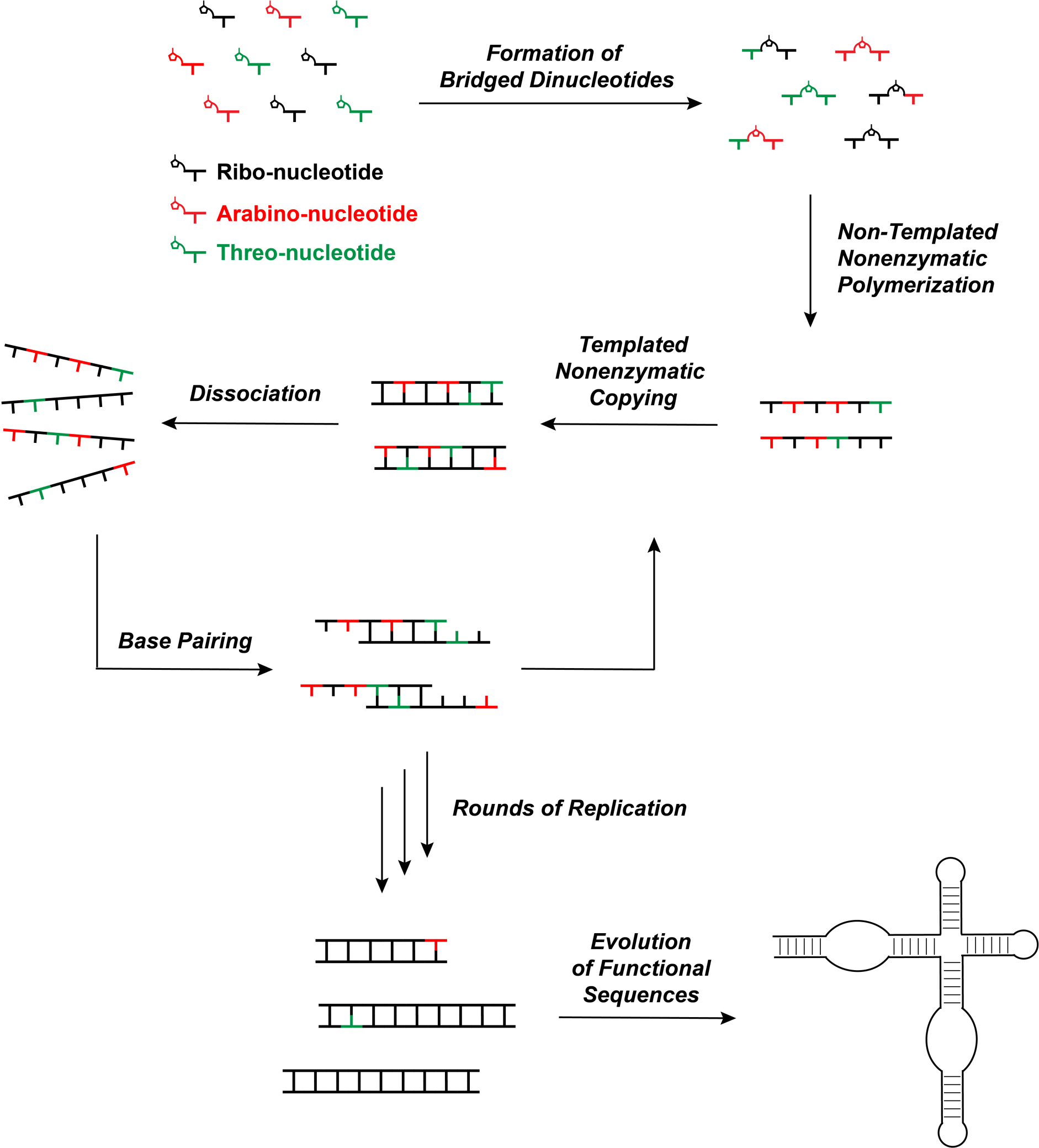
Schematic diagram illustrating pathways for the emergence of RNA from prebiotic mixtures and evolution of functional sequences.

Consistent with nonenzymatic template-directed copying, non-templated primer extension primarily occurs through imidazolium-bridged dinucleotides. We observed a much faster rate of non-templated primer extension with bridged dinucleotides than with activated mononucleotides (Figure 2F, G). The superior leaving group of the bridged dinucleotides makes them the primary substrates in both non-templated and templated primer extension. The actual reaction rates varied with time (Figure 2D, E), because of the time required for formation of bridged dinucleotides from monomers and the more rapid hydrolysis of bridged dinucleotides than activated monomers (Figure S3). In a prebiotic scenario, the extent of non-templated oligomerization would depend strongly on the effectiveness of the ambient activation chemistry, as well as on environmental factors such as pH and temperature that would affect both activation and hydrolysis.

Contrary to individual reactivity assays where threo-nucleotide incorporation is disfavored by a factor of 6 relative to ribonucleotides (Figure 3F), competition experiments show a reduced bias against threo-nucleotides (Figure 4, 6). A possible explanation for this difference is that in a mixture, hetero-bridged dinucleotides such as rN*tN and araN*tN (Table S5) would form more readily than tN*tN, due to the less hindered attack by the 2AI moiety of a *tN on the phosphate groups of *rN or *araN nucleotides. By mitigating the sterically crowded nature of threo-nucleotides, the formation of hetero-bridged dinucleotides may facilitate the incorporation of threo-nucleotides in non-templated reactions. This mechanism is consistent with the observation that activated downstream oligonucleotides accelerate template-directed primer extension with threo-nucleotides (17).

The poor regioselectivity of non-templated polymerization results in significant formation of 2ʹ-5ʹ linkages (33). In addition, internal 2ʹ hydroxyl groups in the primer can react with activated nucleotides forming branched structures (Figure 5). Our competition experiments suggest that sugar heterogeneity will be enhanced by the copolymerization of mixed nucleotides (Figure 4, 6). However, this initial heterogeneity is likely to be reduced during subsequent template-directed copying steps. For example, a template containing a 2ʹ-5ʹ linkage is copied at a reduced rate but no detectable 2ʹ-5ʹ linkages are formed in the product (34). Similarly, arabino- and threo-nucleotides are outcompeted by ribonucleotides in template directed copying experiments (15, 17). On the other hand, a degree of backbone heterogeneity is compatible with RNA folding into functional structures and might also be helpful for strand separation by lowering the melting temperature of product duplexes, thus facilitating subsequent cycles of primer extension (35, 36). 2ʹ-5ʹ branched RNA structures (37) form biologically as the lariat products of RNA splicing (38), suggesting that branched RNAs could also have prebiotic roles. In the presence of complementary oligomers, the probability of reactions at internal 2ʹ-OHs decreases markedly (Figure 5H, Table S4B). While branched products may not be ideal for propagating genetic information, they might still serve as splints to assist template-directed copying (39, 40) and RNA replication under the VCG Model (32).

In summary, our study strengthens our understanding of the physico-chemical selection steps that led to the emergence of RNA from prebiotic mixtures. The copolymerization of mixtures of different types of nucleotides results in extension products that roughly mirror the input ratio of nucleotides. This process results in a modest bias against threo-nucleotide incorporation, although the presence of ribo- and arabino-nucleotides actually mitigates the intrinsic bias against threo-nucleotides. As a result, bias in favor of ribonucleotides during synthesis would be retained during non-templated polymerization, and then more strongly enhanced during subsequent template-directed reactions (15, 17). The combined selection in favor of ribonucleotides across sequential physical and chemical steps likely resulted in RNA-enriched oligomers after successive rounds of replication, setting the stage for the evolution of complex, functional RNA sequences (Figure 7).

## DATA AVAILABILITY

The data underlying this article are available in the article and in its online supplementary material.

## SUPPLEMENTARY DATA

Supplementary information are available online.

## ACKNOWLEDGEMENTS

The authors thank Professor John C. Chaput for providing the threo-nucleosides. We thank Dr. Victor Lelyveld, Dr. Harry Aitken, and Dr. Seohyun Chris Kim for helpful discussions and technical assistance. We thank Dr. Victor Lelyveld, Dr. Saurja DasGupta, and Dr. Filip Bošković for comments on the manuscript.

