## Supplementary Information for "Constraints on the Emergence of RNA through Non-Templated Primer Extension with Mixtures of Potentially Prebiotic Nucleotides"

**1 Materials and Methods**
**1.1 Characterization of Activated Mononucleotides and Bridged Dinucleotides**
1.1.1 ribo-cytidine-5'-phosphoro-(2-aminoimidazole) **(*rC)**
1.1.2 $^{13}$C, $^{15}$N-ribo-cytidine-5'-phosphoro-(2-aminoimidazole) **(isotope-labeled *rC)**
1.1.3 arabino-cytidine-5'-phosphoro-(2-aminoimidazole) **(*araC)**
1.1.4 threo-cytidine-5'-phosphoro-(2-aminoimidazole) **(*tC)**
1.1.5 1,3-di-(cytidine-5'-phosphoryl)-2-aminoimidazolium **(rC*rC)**
1.1.6 ribo-guanosine-5'-phosphoro-(2-aminoimidazole) **(*rG)**
1.1.7 arabino-guanosine-5'-phosphoro-(2-aminoimidazole) **(*araG)**
1.1.8 threo-guanosine-5'-phosphoro-(2-aminoimidazole) **(*tG)**
1.1.9 1,3-di-(guanosine-5'-phosphoryl)-2-aminoimidazolium **(rG*rG)**
**1.2 Oligonucleotide Synthesis**
**2 Supplementary Scheme and Figures**
**3 Supplementary Tables**

### 1 Materials and Methods

#### 1.1 Characterization of Activated Mononucleotides and Bridged Dinucleotides

##### 1.1.1 ribo-cytidine-5'-phosphoro-(2-aminoimidazole) (*rC)

**$^{1}$H NMR** (400 MHz, $D_2O$) δ 7.75 (d, *J* = 7.6 Hz, 1H), 6.80 (m, 1H), 6.61 (m, 1H), 6.05 (d, *J* = 7.6 Hz, 1H), 5.95 (d, *J* = 4.2 Hz, 1H), 4.25 (t, *J* = 4.5 Hz, 1H), 4.24 – 4.16 (m, 3H), 4.09 (m, 1H).
**$^{31}$P NMR** (162 MHz, $D_2O$) δ -10.74.
**HRMS** (Q-TOF) m/z: calculated for $[C_{12}H_{16}N_6O_7P]^-$ 387.0896; found: 387.0860.

##### 1.1.2 $^{13}$C, $^{15}$N-ribo-cytidine-5'-phosphoro-(2-aminoimidazole) (isotope-labeled *rC)

**$^{1}$H NMR** (400 MHz, $D_2O$) δ 7.57 (d, 1H), 6.49 – 6.43 (m, 2H), 5.85 (d, 1H), 5.64 (s, 1H), 4.62 (dd, *J* = 8.6 Hz, 2H), 4.39 – 4.27 (m, 2H), 3.94 (s, 1H).
**$^{31}$P NMR** (162 MHz, $D_2O$) δ -9.57.
**HRMS** (Q-TOF) m/z: calculated for $[C_3{}^{13}C_9H_{16}N_3{}^{15}N_3O_7P]^-$ 399.1109; found: 399.1112.

##### 1.1.3 arabino-cytidine-5'-phosphoro-(2-aminoimidazole) (*araC)

**$^{1}$H NMR** (400 MHz, $D_2O$) δ 7.63 (d, *J* = 7.6 Hz, 1H), 6.81 (s, 1H), 6.61 (s, 1H), 6.20 (s, 1H), 6.03 (d, *J* = 7.4 Hz, 1H), 4.40 (m, 1H), 4.23 – 4.01 (m, 4H).
**$^{31}$P NMR** (162 MHz, $D_2O$) δ -10.68.
**HRMS** (Q-TOF) m/z: calculated for $[C_{12}H_{16}N_6O_7P]^-$ 387.0896; found: 387.0885.

##### 1.1.4 threo-cytidine-5'-phosphoro-(2-aminoimidazole) (*tC)

**$^{1}$H NMR** (400 MHz, $D_2O$) δ 7.66 (d, *J* = 7.6 Hz, 1H), 6.58 (td, *J* = 19.3 Hz, 2H), 5.93 (d, *J* = 7.6 Hz, 1H), 5.72 (s, 1H), 4.73 (dd, *J* = 8.3, 3.2 Hz, 1H), 4.49 – 4.37 (m, 2H), 4.06 (s, 1H).
**$^{31}$P NMR** (162 MHz, $D_2O$) δ -13.16.
**HRMS** (Q-TOF) m/z: calculated for $[C_{11}H_{14}N_6O_6P]^-$ 357.0791; found: 357.0787.

##### 1.1.5 1,3-di-(cytidine-5'-phosphoryl)-2-aminoimidazolium (rC*rC)

**$^{1}$H NMR** (400 MHz, $D_2O$) δ 7.63 (t, *J* = 7.7 Hz, 2H), 6.93 (t, *J* = 2.0 Hz, 2H), 6.01 (d, *J* = 7.6 Hz, 2H), 5.82 (d, *J* = 3.2 Hz, 2H), 4.28 – 4.09 (m, 10H).
**$^{31}$P NMR** (162 MHz, $D_2O$) δ -12.84.
**HRMS** (Q-TOF) m/z: calculated for $[C_{21}H_{28}N_9O_{14}P_2]^-$ 692.1387; found: 692.1310.

##### 1.1.6 ribo-guanosine-5'-phosphoro-(2-aminoimidazole) (*rG)

**$^{1}$H NMR** (400 MHz, $D_2O$) δ 7.89 (s, 1H), 6.66 (t, *J* = 0.5 Hz, 1H), 6.46 (t, *J* = 1.8 Hz, 1H), 5.87 (s, 1H), 4.70 (t, *J* = 5.6 Hz, 1H), 4.38 – 4.33 (m, 1H), 4.28 – 4.22 (m, 1H), 4.11 – 4.01 (m, 2H).
**$^{31}$P NMR** (162 MHz, $D_2O$) δ -10.66.
**HRMS** (Q-TOF) m/z: calculated for $[C_{13}H_{16}N_8O_7P]^-$ 427.0958; found: 427.0965.

##### 1.1.7 arabino-guanosine-5'-phosphoro-(2-aminoimidazole) (*araG)

**$^{1}$H NMR** (400 MHz, $D_2O$) δ 7.79 (s, 1H), 6.67 (dd, *J* = 1.9, 1.3 Hz, 1H), 6.46 (t, *J* = 2.1 Hz, 1H), 6.16 (d, *J* = 5.8 Hz, 1H), 4.48 (t, *J* = 5.9 Hz, 1H), 4.33 (t, *J* = 6.2 Hz, 1H), 4.18 – 3.99 (m, 3H).
**$^{31}$P NMR** (162 MHz, $D_2O$) δ -10.59.
**HRMS** (Q-TOF) m/z: calculated for $[C_{13}H_{16}N_8O_7P]^-$ 427.0958; found: 427.0966.

##### 1.1.8 threo-guanosine-5'-phosphoro-(2-aminoimidazole) (*tG)

**$^1$H NMR** (400 MHz, $D_2O$) δ 7.64 (d, *J* = 7.6 Hz, 1H), 6.62 – 6.51 (m, 2H), 5.91 (d, *J* = 7.6 Hz, 1H), 5.69 (s, 1H), 4.71 (dd, *J* = 8.3, 3.2 Hz, 1H), 4.47 – 4.34 (m, 1H), 4.04 (s, 1H).
**$^{31}$P NMR** (162 MHz, $D_2O$) δ -12.62.
**HRMS** (Q-TOF) m/z: calculated for $[C_{12}H_{14}N_8O_6P]^-$ 397.0852; found: 397.0862.

1.1.9 1,3-di-(guanosine-5'-phosphoryl)-2-aminoimidazolium **(rG*rG)**

**$^1$H NMR** (400 MHz, $D_2O$) δ 7.83 (s, 2H), 6.71 – 6.66 (m, 2H), 5.75 (d, *J* = 5.2 Hz, 2H), 4.65 (t, *J* = 5.3 Hz, 2H), 4.39 (t, 2H), 4.15 – 4.05 (m, 4H), 4.05 – 3.95 (m, 2H).
**$^{31}$P NMR** (162 MHz, $D_2O$) δ -12.92.
**HRMS** (Q-TOF) m/z: calculated for $[C_{23}H_{28}N_{13}O_{14}P_2]^-$ 772.1354; found: 772.1429.

#### 1.2 Oligonucleotide Synthesis

Oligonucleotides were either purchased from IDT or synthesized in-house by solid phase synthesis on the Expedite 8909 DNA/RNA synthesizer. Phosphoramidites and reagents were purchased from ChemGenes (Wilmington, MA) and Glen Research (Sterling, MA). The synthesized oligonucleotides were deprotected and purified by Glen-Pak RNA purification cartridges (Sterling, MA).

#### 2 Supplementary Scheme and Figures

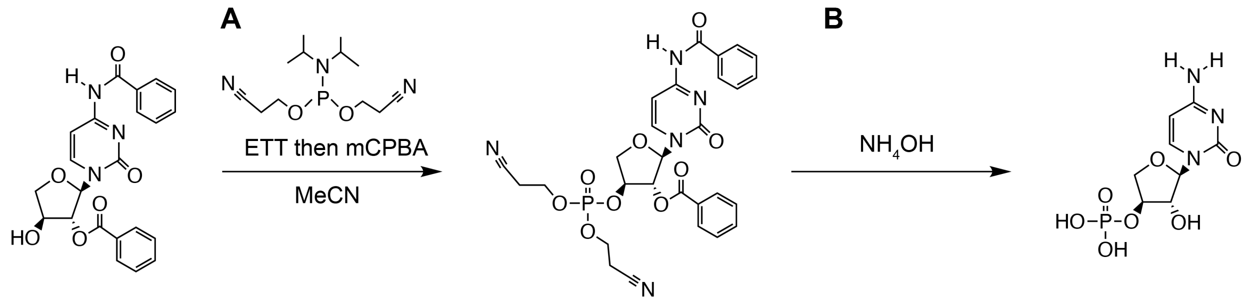


**TNA C nucleoside** **tCMP**

**Scheme S1.** Synthesis of threose cytidine monophosphate (tCMP) from (A) phosphorylation and (B) deprotection of TNA C nucleoside.

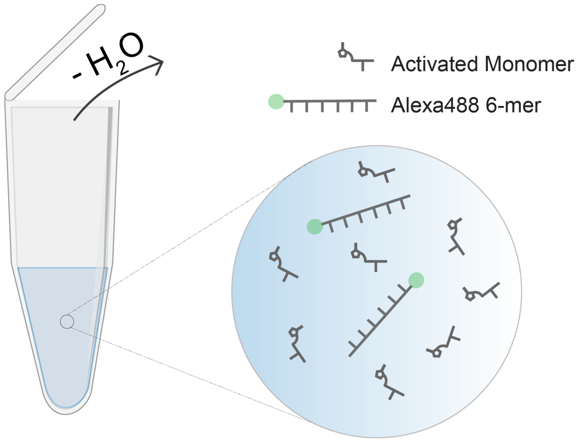


**Figure S1.** Schematic representation of the spontaneous air-drying experiments. Water in the reaction mixture gradually evaporates under ambient air flow and dry down to a clear paste at the bottom of the Eppendorf tube. During this process, both the primer (XJ-Alexa-6mer, as listed in Table S1) and the activated mononucleotides become concentrated. As a result, the non-templated primer extension speeds up to generate considerable amount of extended products.

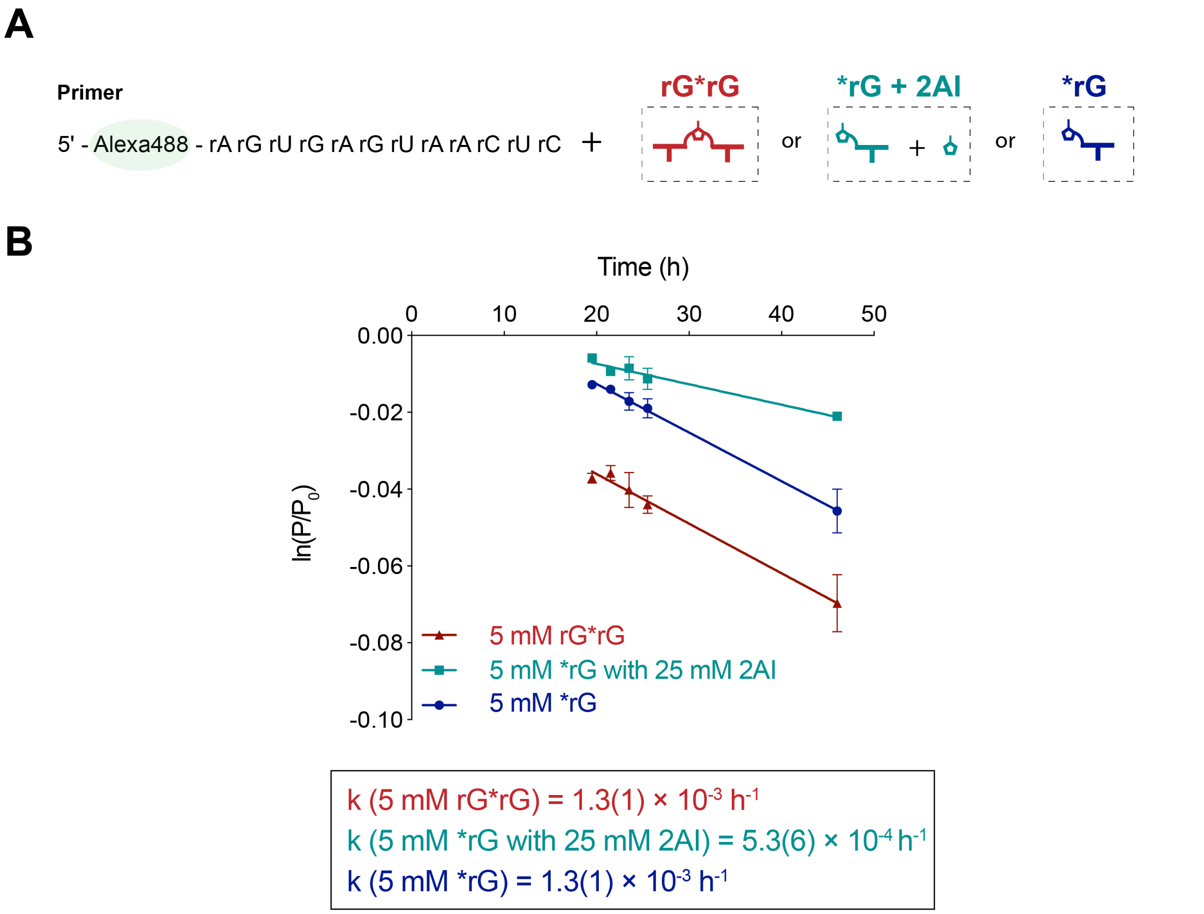


**Figure S2.** Mechanism of non-templated primer extension. (A) Schematic representation of a non-templated primer extension reaction using an Alexa488-labeled RNA 12-mer with bridged dinucleotides (rG*rG) or activated mononucleotides (*rG). 2AI was added to reduce the formation of bridged dinucleotides. (B) Gel electrophoresis images of non-templated primer extension reactions using bridged dinucleotides or activated mononucleotides. (C) Kinetic analysis with observed pseudo-first-order reaction rates ($k_{obs}$). Reaction conditions: 1 μM primer (XJ-Alexa-12mer, Table S1), 200 mM HEPES at pH 8.0, and 50 mM $MgCl_2$ with 5 mM rG*rG, 5 mM *rG with 25 mM 2AI, or 5 mM *rG. Error bars indicate standard deviations of the mean, n=2 replicates.

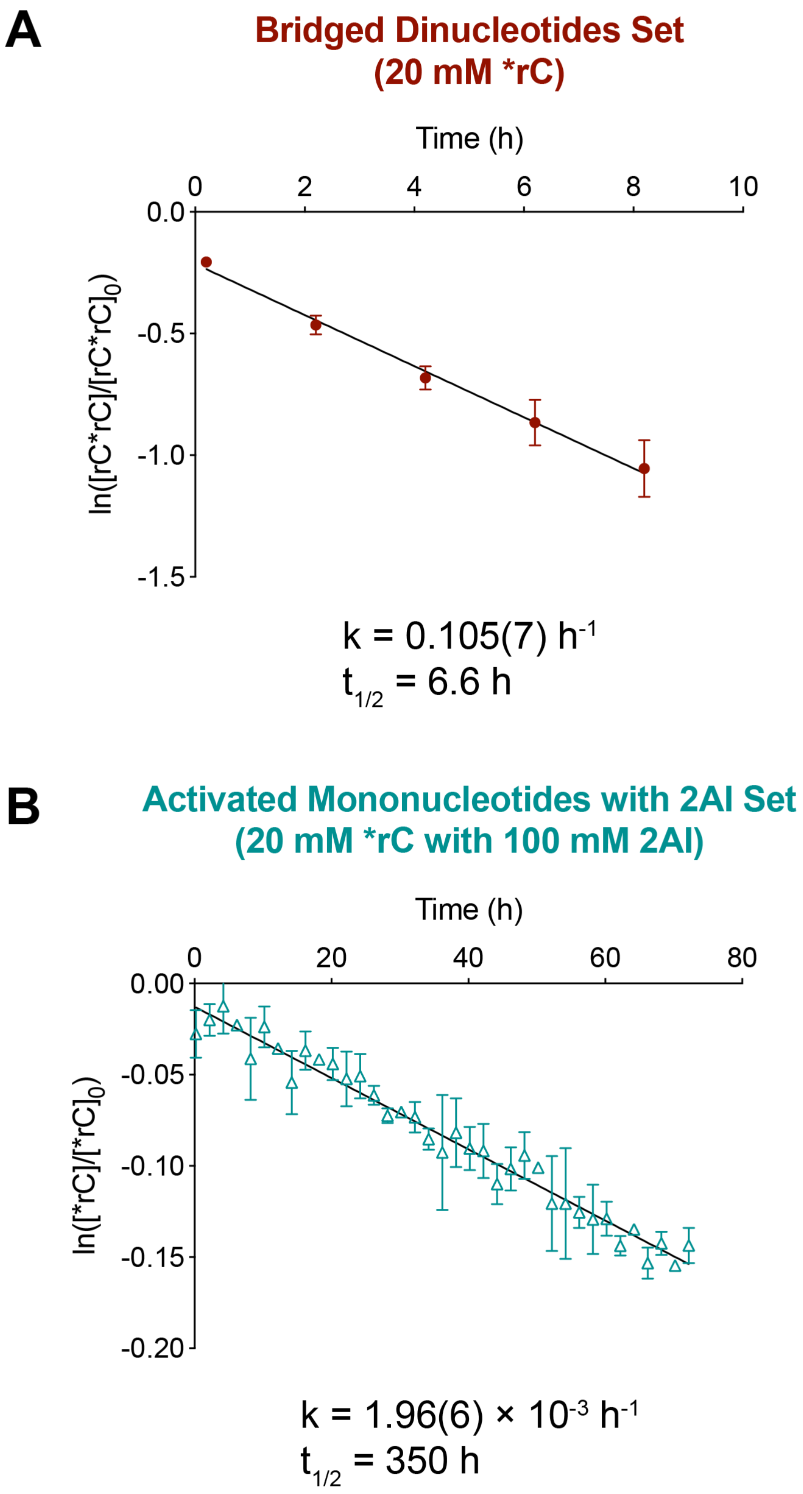


**Figure S3.** The hydrolysis rates and half-lives of (A) bridged dinucleotides (20 mM rC*rC) and (B) activated mononucleotides with 2AI (20 mM *rC with 100 mM 2AI) derived from the $^{31}$P NMR experiments in Figure S5. Reaction conditions: the respective concentration of activated species as indicated above, 50 mM $MgCl_2$, 200 mM HEPES at pH 8, 10% (v/v) $D_2O$, incubation at RT. Error bars indicate standard deviations of the mean, n=2 replicates.

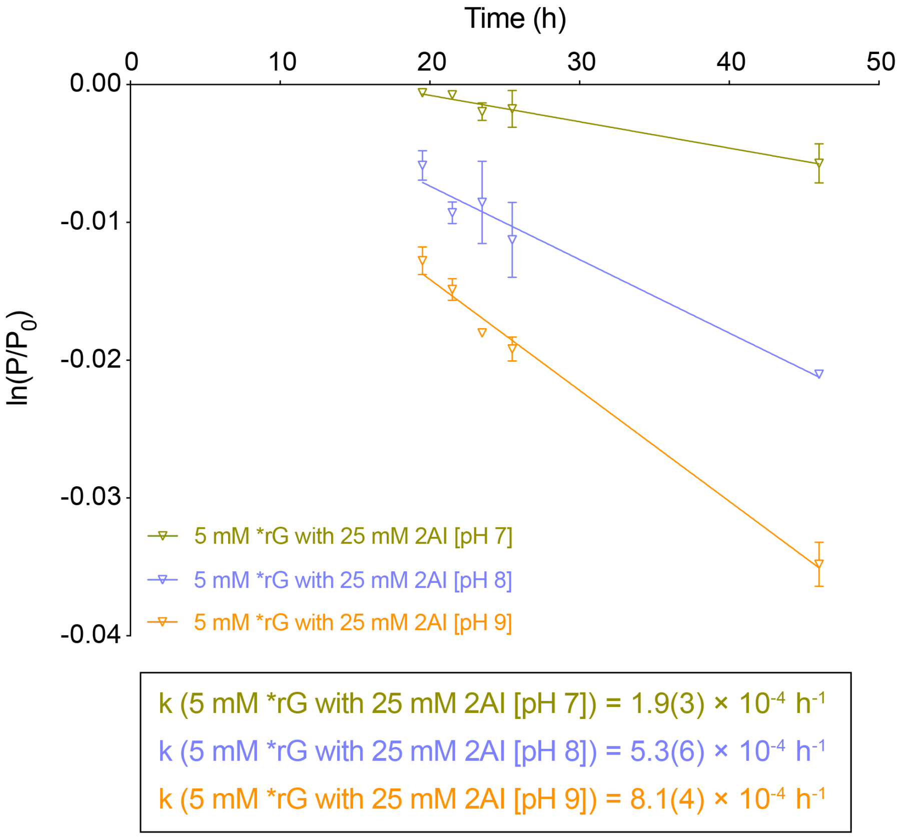


**Figure S4.** Non-templated primer extension with activated monomers with 2AI at different pHs. Reaction conditions: 5 mM *rG with 5-fold 2AI or 5 mM *rG were added to 1 μM primer (XJ-Alexa-12mer, Table S1),, 200 mM HEPES pH 7 or 8 or 9 and 50 mM $MgCl_2$ respectively. Error bars indicate standard deviations of the mean, n=2 replicates.

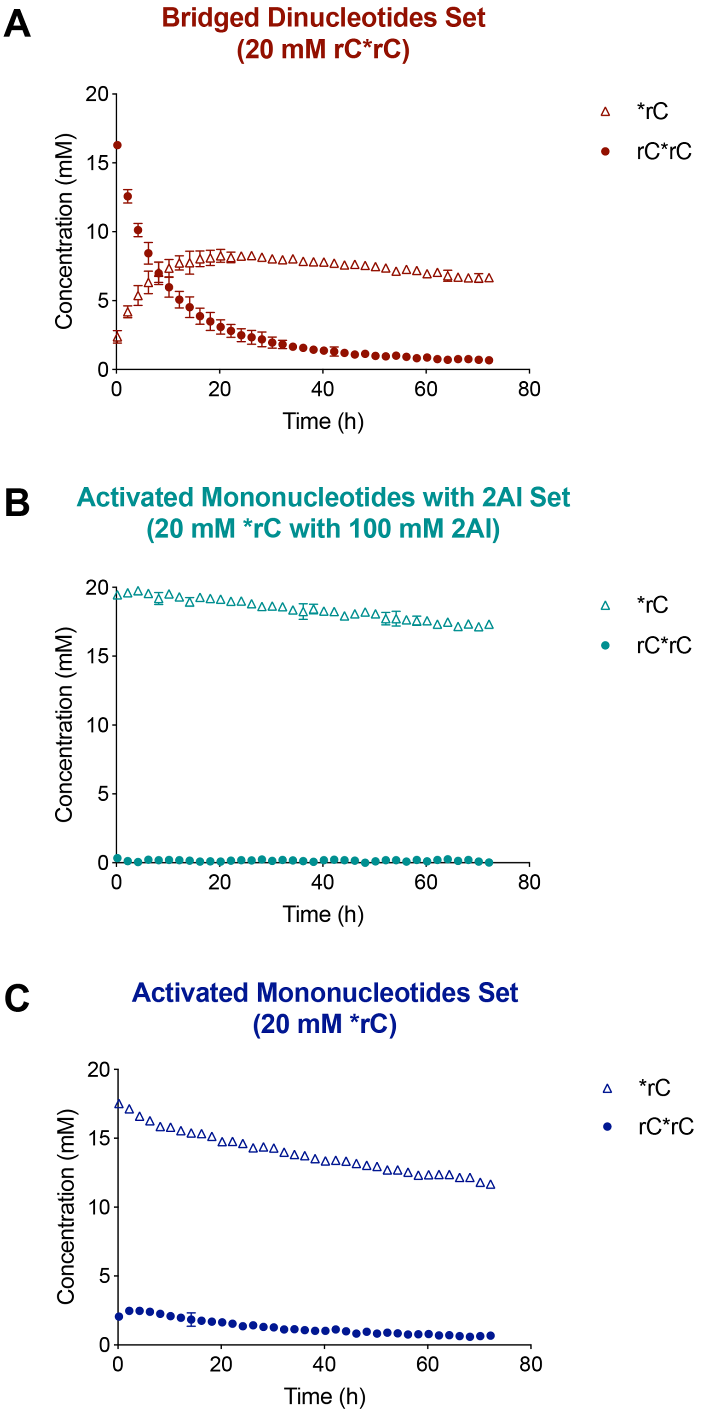


**Figure S5.** $^{31}P$ NMR hydrolysis experiments of (A) bridged dinucleotides (20 mM rC*rC), (B) activated mononucleotides with 2AI (20 mM *rC with 100 mM 2AI), and (C) activated mononucleotides alone sets (20 mM *rC). Plot of the concentration of active species (*rC and rC*rC) in solution over time. Reaction conditions: the respective concentration of activated species as indicated above, 50 mM $MgCl_2$, 200 mM HEPES at pH 8, 10% (v/v) $D_2O$, incubation at RT. Error bars indicate standard deviations of the mean, n=2 replicates.

**A**

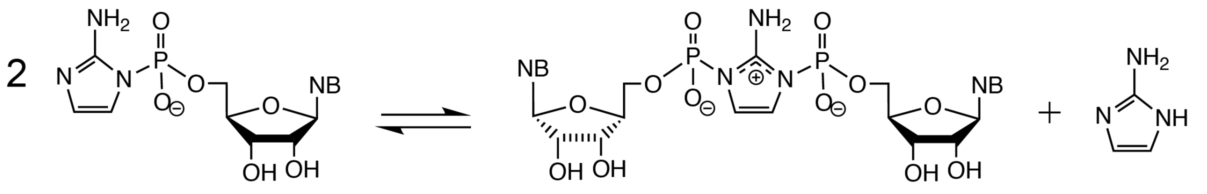

***rN** **rN*rN** **2AI**

**B**

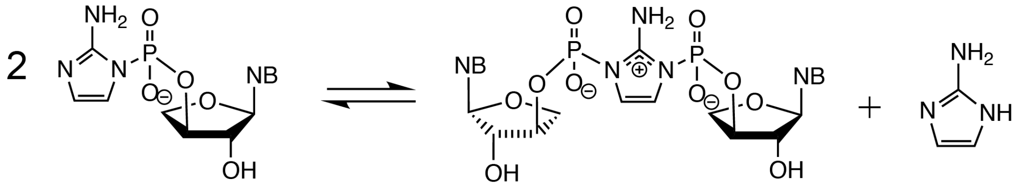

***tN** **tN*tN** **2AI**

**Figure S6.** Schematic representation of the formation of (A) 5'-5' imidazolium-bridged ribonucleotides (rN*rN) and (B) 3'-3' imidazolium-bridged threo-nucleotides (tN*tN).

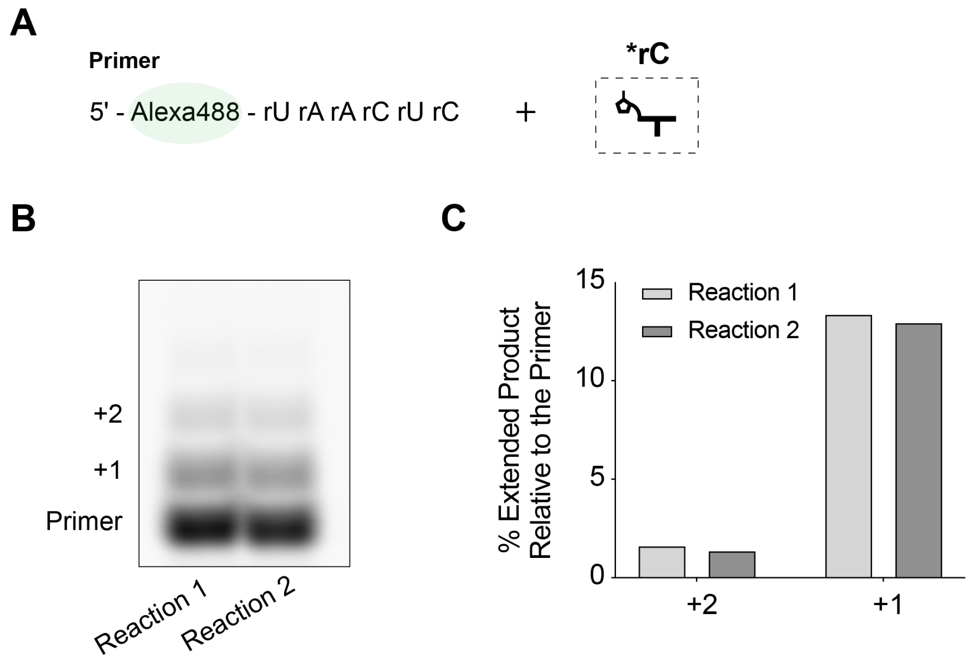


**Figure S7.** Variability of spontaneous air-drying experiments within the same set (performed at the same time and exposed to the identical ambient air flow rate and moisture level). (A) Schematic representation of non-templated primer extension reaction using an Alexa488-labeled RNA 6-mer with activated mononucleotides (*rC). (B) Gel electrophoresis image of non-templated primer extension reactions within the set. (C) Quantification of extended products relative to the primer (100%). Reaction conditions: 10 μM primer (XJ-Alexa-6mer, Table S1), 200 mM HEPES pH at 8.0, 50 mM $MgCl_2$, and 20 mM *rC. The reaction mixtures were subjected to air-drying to clear pastes under ambient air to accelerate non-templated primer extension (as previously described in Figure S1) and were subsequently quenched at 24h.

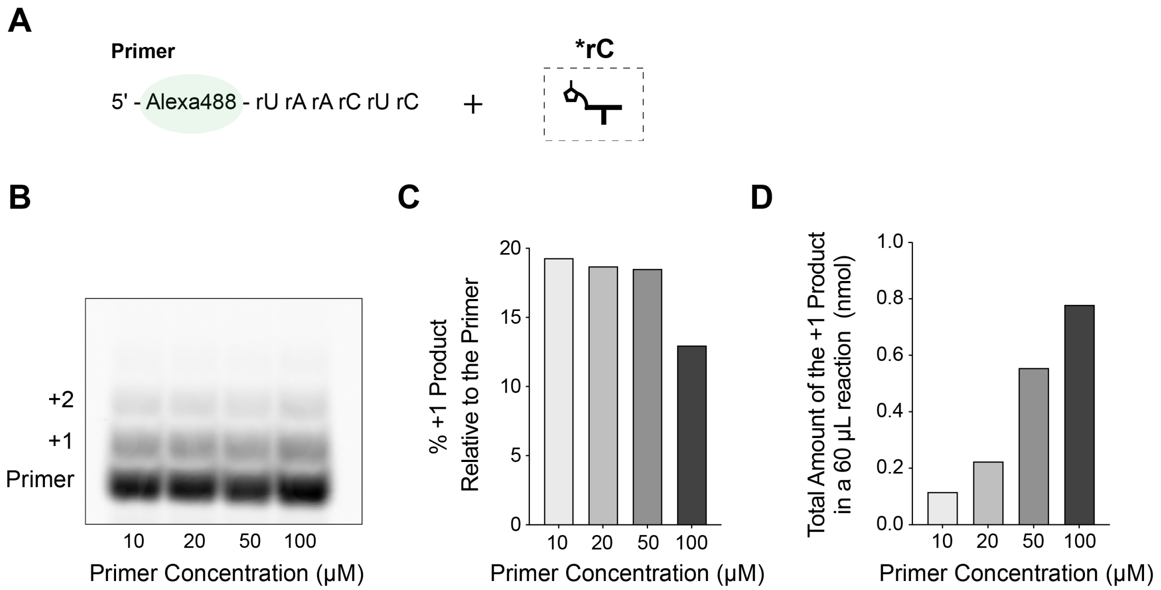


**Figure S8.** Effect of primer concentrations on spontaneous air-drying experiments. (A) Schematic representation of non-templated primer extension reaction using an Alexa488 labeled RNA 6-mer (Table S1) with activated mononucleotides (*rC). (B) Gel electrophoresis image of non-templated primer extension reactions across different primer concentrations. (C) Quantification of +1 product relative to primer (100%). (D) Derived total amount of +1 product at different primer concentrations for a 60 μL reaction. 100 μM primer concentration yielded the highest amount of +1 product for characterization and was therefore chosen as the most optimal condition for subsequent spontaneous air-drying and competition experiments. Reaction conditions: primer (10 μM, 20 μM, 50 μM or 100 μM), 200 mM HEPES pH 8.0, 50 mM $MgCl_2$ and 20 mM *rC. The reaction mixtures were dried down to clear pastes under ambient air to speed up non-templated primer extension (Figure S1) and were subsequently quenched at 24h.

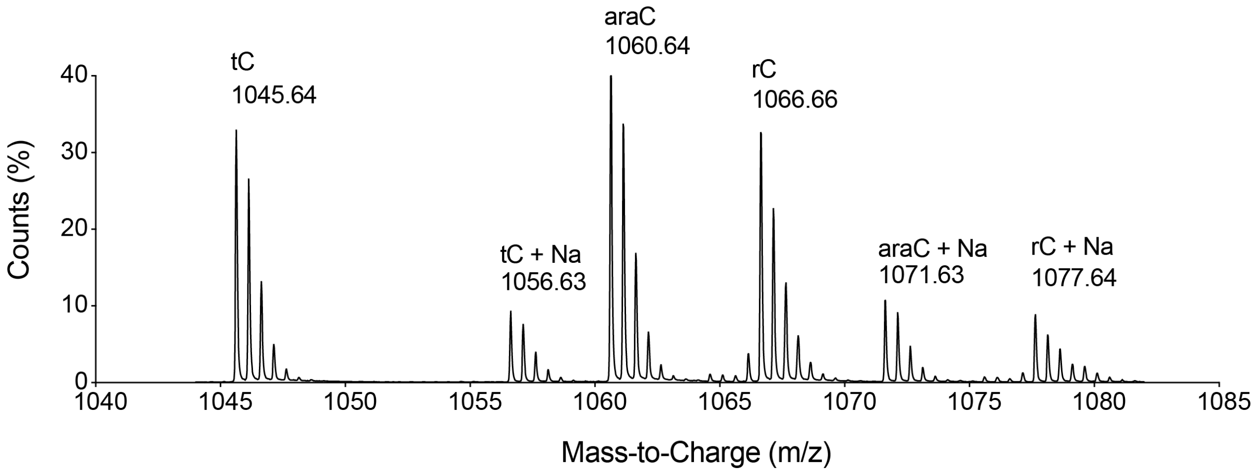


**Figure S9.** Plot of counts (%) as a function of mass-to-charge (m/z) of the +1 products and their salt adducts of *rC:*araC:*tC = 1:1:1 competition experiment using an 5'-OH RNA 6-mer. Species shown are in the -2 charged state $(M-2H)^{2-}$. Reaction conditions: 100 μM 6-mer primer (XJ-5, Table S1) and 6.7 mM each of *rC, *araC and *tC. The reactions were allowed to dry spontaneously under ambient air and were subsequently quenched at 24 h. The samples were then desalted on C18 ZipTip pipette tips, followed by injection in HR LC-MS. The generated compound lists using the Agilent MassHunter Qualitative Analysis software were matched with the calculated masses of all possible +1 non-templated primer extension products and their salt adducts (Table S2).

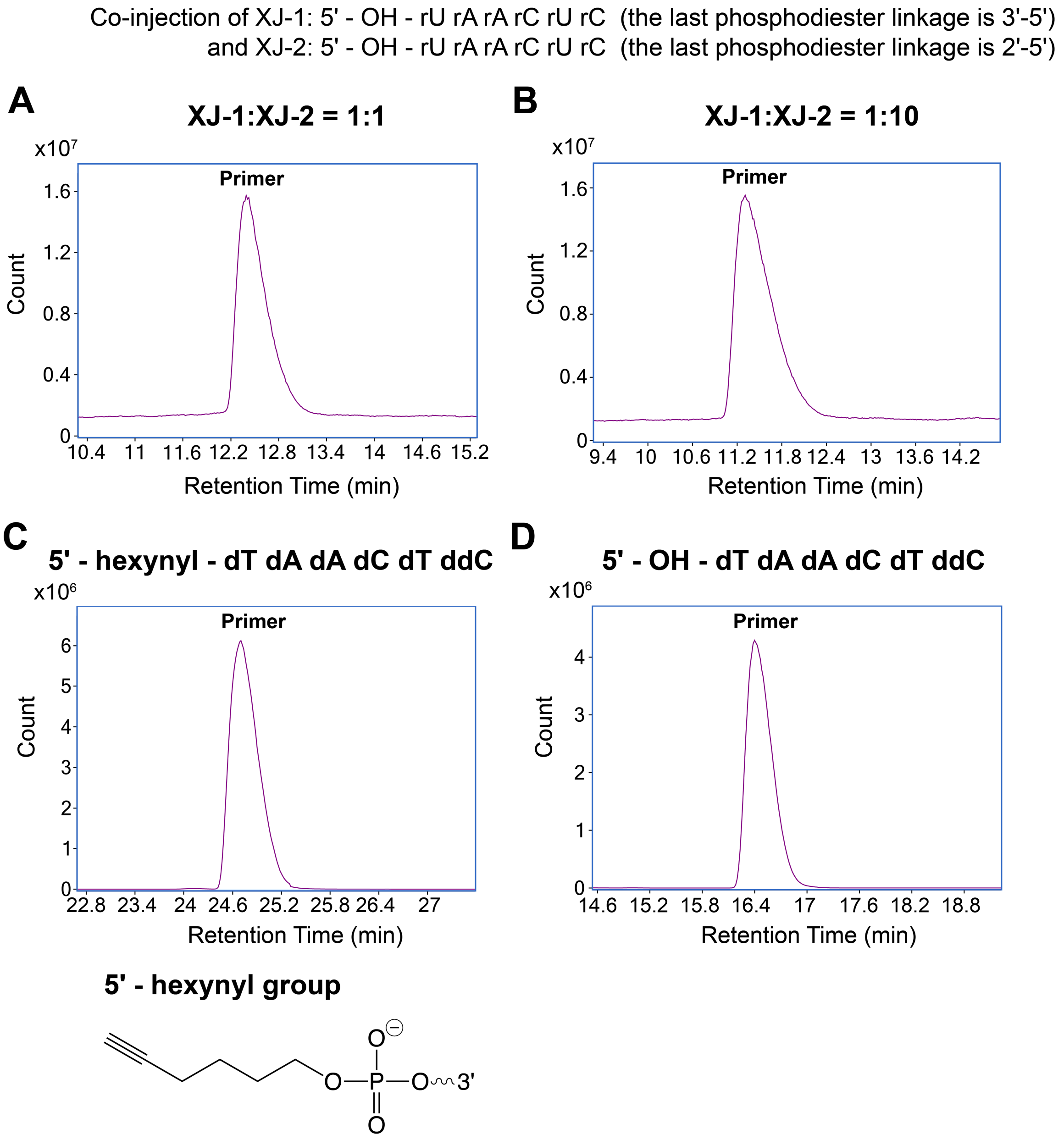


**Figure S10.** Origin of the double peak pattern in the chromatograms of the competition experiments in Figure 4. (A-B) Total ion chromatograms (TIC) from the co-injection of oligomers ending with 3'-5' (XJ-1, Table S1) and 2'-5' phosphodiester linkages (XJ-2, Table S1) in (A) 1:1 ratio of XJ-1 and XJ-2 and (B) 1:10 ratio of XJ-1 and XJ-2. Note that the XJ-1 and XJ-2 oligomers were synthesized using TOM-protected RNA amidites to exclude the possibility of 2' to 3' migration of the TBDMS group. Standards with 3'-5' and 2'-5' linkage could not be separated on LC-MS. (C) TIC of the non-templated primer extension product of a 5'-hexynyl DNA primer (XJ-9, Table S1) showed that nucleobases did not react. The chemical structure of the 5'-hexynyl group is indicated. (D) TIC of the non-templated primer extension product of a 5'-OH DNA primer (XJ-11, Table S1) showed that 5'-OH did not react with the activated mononucleotides. Reaction conditions of (C)-(D): 20 mM activated mononucleotides, 100 μM 6-mer primer, 200 mM HEPES at pH 8.0, and 50 mM $MgCl_2$. The reactions were allowed to dry spontaneously under ambient air and were subsequently quenched at 24 h. The samples were then desalted on C18 ZipTip pipette tips, followed by injection in HR LC-MS.

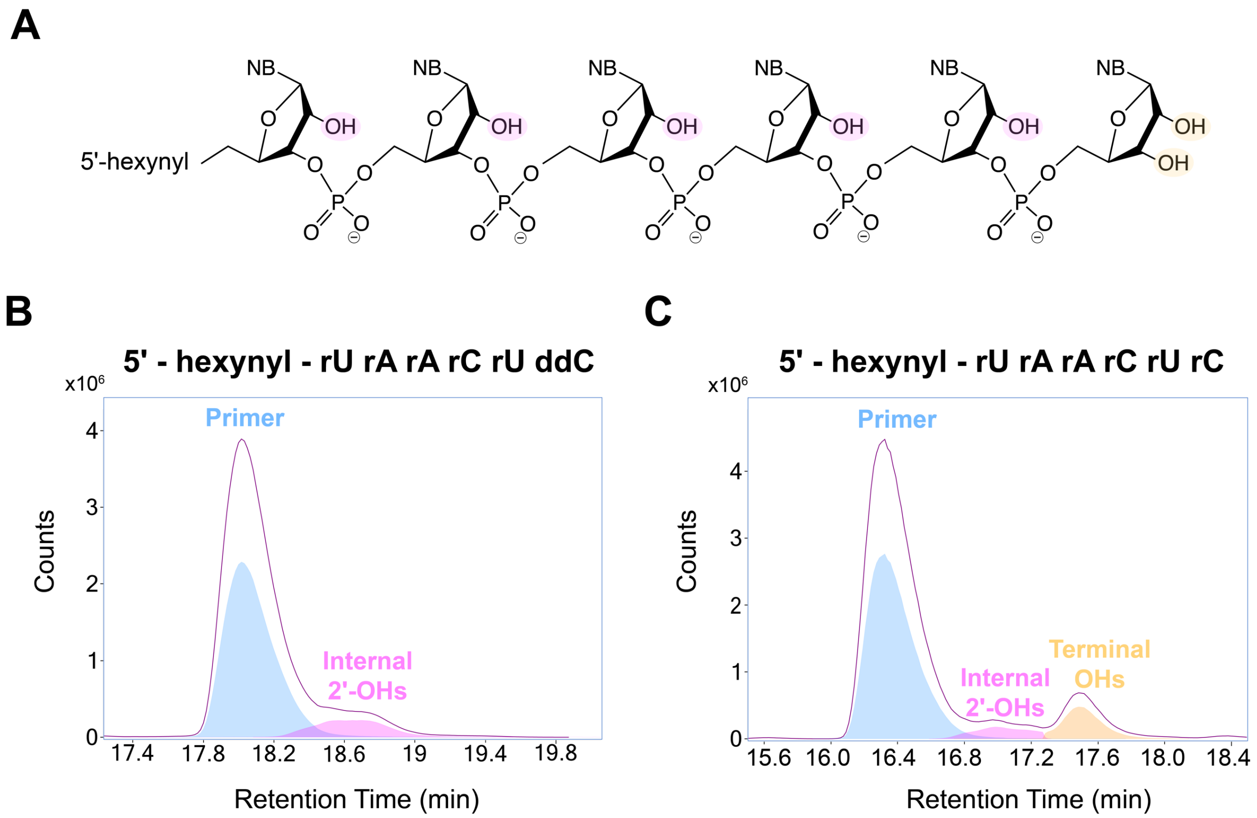


**Figure S11.** Both internal and terminal hydroxyls can react in non-templated primer extension. (A) Schematic representation of the internal 2'-OHs and terminal OHs in an RNA oligomer (5'-hexynyl-rU rA rA rC rU rC). Overlay of TCC (purple) and ECC (blue: primer; pink: internal 2'-OH reaction; yellow: terminal OH extension) of the non-templated primer extension product with (B) 5'-hexynyl RNA primer ending with dideoxy cytosine (XJ-7, Table S1) and (C) 5'-hexynyl RNA primer (XJ-8, Table S1). Reaction conditions for (B) and (C): 20 mM activated mononucleotides, 100 μM 6-mer primer, 200 mM HEPES at pH 8.0, and 50 mM $MgCl_2$. The reactions were allowed to dry spontaneously under ambient air and were subsequently quenched at 24 h. The samples were then desalted on C18 ZipTip pipette tips, followed by injection in HR LC-MS.

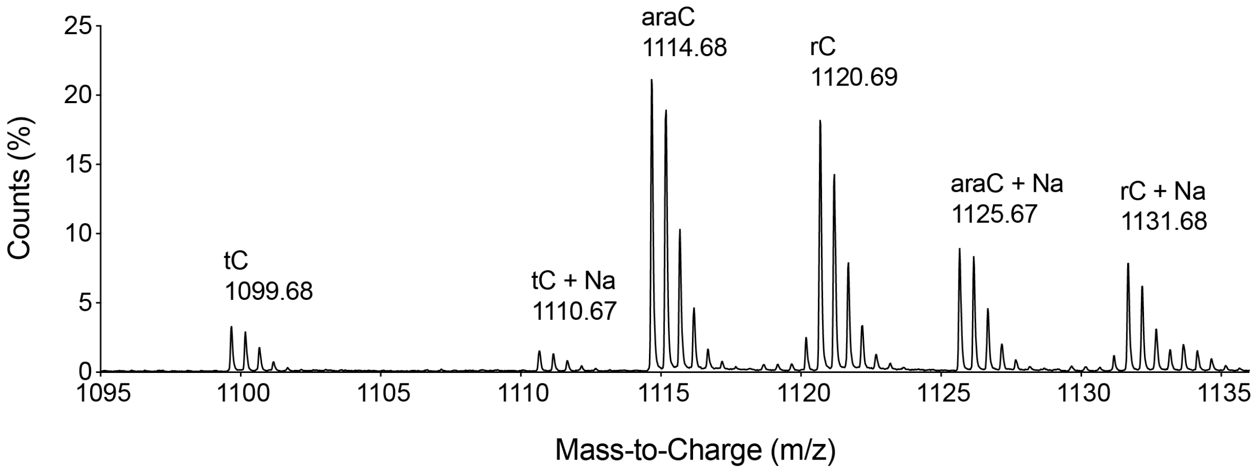


**Figure S12.** Ion counts (%) as a function of mass-to-charge (m/z) of the +1 products and their salt adducts of *rC:*araC:*tC = 1:1:1 competition experiment (Reaction1, Set 3) with a modified primer. All mass-to-charge ratios here are in the -2 charged state $(M-2H)^{2-}$. Reaction conditions: 100 μM 6-mer primer (XJ-10, Table S1) and 6.7 mM each of *rC, *araC and *tC. The reactions were allowed to dry spontaneously under ambient air and were subsequently quenched at 24 h. The samples were then desalted on C18 ZipTip pipette tips, followed by injection in HR LC-MS. The generated compound lists using the Agilent MassHunter Qualitative Analysis software were matched with the calculated masses of all possible +1 non-templated primer extension products and their salt adducts (Table S3). For further information on the HR LC-MS, please refer to the materials and methods section.

#### 3 Supplementary Tables

**Table S1. Sequences of oligonucleotides used in the study.**

| Name | Source | Sequence (5' → 3') |
|---|---|---|
| XJ-FAM-12mer | IDT | 5' - FAM - rA rG rU rG rA rG rU rA rA rC rU rC |
| XJ-Alexa-6mer | IDT | 5' - Alexa 488 - rU rA rA rC rU rC |
| XJ-Alexa-12mer | IDT | 5' - Alexa 488 - rA rG rU rG rA rG rU rA rA rC rU rC |
| XJ-1 | In-house | 5' - OH - rU rA rA rC rU rC rC (last linkage: 3'- 5') |
| XJ-2 | In-house | 5' - OH - rU rA rA rC rU rC rC (last linkage: 2'- 5') |
| XJ-5 | IDT | 5' - OH - rU rA rA rC rU rC |
| XJ-7 | IDT | 5' - hexynyl - rU rA rA rC rU ddC |
| XJ-8 | IDT | 5' - hexynyl - rU rA rA rC rU rC |
| XJ-9 | IDT | 5' - hexynyl - dT dA dA dC dT ddC |
| XJ-10 | IDT | 5' - hexynyl - dT dA dA dC dT rC |
| XJ-11 | IDT | 5' - OH - dT dA dA dC dT ddC |
| XJ-6mer-rc | IDT | 5' - OH - rG rA rG rU rU rA |

**Table S2. Observed and calculated masses of the expected products of competition experiments.**

| | Observed Mass (Da) | Calculated Mass (Da) |
|---|---|---|
| **Primer + rC** | 2135.33 | 2135.35 |
| **Primer + araC** | 2123.31 | 2123.32 |
| **Primer + tC** | 2093.30 | 2093.31 |
| **Primer + rC + rC** | 2452.39 | 2452.42 |
| **Primer + araC + araC** | 2428.34 | 2428.36 |
| **Primer + tC + tC** | N/A | 2368.34 |
| **Primer + rC + araC *** | 2440.37 | 2440.39 |
| **Primer + rC + tC *** | 2410.35 | 2410.38 |
| **Primer + araC + tC *** | 2398.33 | 2398.35 |

* Sequence might not be in order.

**Table S3. Observed and calculated masses of the expected products of competition experiments with a 5'-hexynyl DNA primer with a 3'-terminal ribonucleotide.**

| | Observed Mass (Da) | Calculated Mass (Da) |
|---|---|---|
| **Primer + rC** | 2243.42 | 2243.43 |
| **Primer + araC** | 2231.40 | 2231.41 |
| **Primer + tC** | 2201.38 | 2201.40 |

**Table S4. Sugar and backbone composition of non-templated products. (A) Nucleotide incorporation (%) of the +1 extended product in the *rC:*araC:*tC = 1:1:1 and 10:1:1 competition experiments. (B) Total and individual extension ratio (internal: terminal) in the non-templated primer extension of 5'-hexynyl RNA primer without and with complementary oligomers. (C) Nucleotide incorporation (%) of the +1 extended product in *rC:*araC:*tC = 1:1:1 and 10:1:1 competition experiments with a modified primer. When calculating the incorporation percentage, the integrated intensity is taken into account from both the +1 extended products and their salt adducts.** [a]

**A.** ***Incorporation of ribo-, arabino-, and threo-nucleotides in competition experiments*** [b]

| *rC:*araC:*tC = 1:1:1 | rC (%) | araC (%) | tC (%) |
|---|---|---|---|
| Reaction 1, Set 1 | 30.6 | 47.1 | 22.3 |
| Reaction 2, Set 1 | 29.5 | 48.4 | 22.2 |
| Reaction 1, Set 2 | 31.8 | 41.9 | 26.3 |
| Reaction 2, Set 2 | 31.7 | 41.5 | 26.8 |
| *rC:*araC:*tC = 10:1:1 | rC (%) | araC (%) | tC (%) |
| Reaction 1, Set 2 | 86.7 | 8.7 | 4.6 |
| Reaction 2, Set 2 | 84.9 | 8.1 | 6.9 |

**B.** ***Presence of complementary oligomers disfavoring extension through internal hydroxyls*** [c]

| Without complementary oligomers | Total Extension Ratio Internal : Terminal | Average Individual Extension Ratio [d] Internal : Terminal |
|---|---|---|
| Reaction 1, Set 1 | 1.0 : 1.6 | 1.0 : 4.0 |
| Reaction 2, Set 1 | 1.0 : 1.5 | 1.0 : 3.8 |
| With complementary oligomers | Total Extension Ratio Internal : Terminal | Average Individual Extension Ratio [d] Internal : Terminal |
| Reaction 1, Set 1 | 1.0 : 4.0 | 1.0 : 10.0 |
| Reaction 2, Set 1 | 1.0 : 3.6 | 1.0 : 9.0 |
| Reaction 3, Set 1 | 1.0 : 3.4 | 1.0 : 8.4 |

**C.** ***Competition experiments with a 5'-hexynyl DNA primer with a 3'-terminal ribonucleotide***

| *rC:*araC:*tC = 1:1:1 | rC (%) | araC (%) | tC (%) |
|---|---|---|---|
| Reaction 1, Set 1 | 41.1 | 43.7 | 15.3 |
| Reaction 2, Set 1 | 40.8 | 43.4 | 15.8 |
| Reaction 1, Set 2 | 37.8 | 44.7 | 17.5 |
| Reaction 2, Set 2 | 37.6 | 42.5 | 20.0 |
| Reaction 1, Set 3 | 38.5 | 47.4 | 14.1 |
| Reaction 2, Set 3 | 38.0 | 48.1 | 14.0 |
| *rC:*araC:*tC = 10:1:1 | rC (%) | araC (%) | tC (%) |
| Reaction 1, Set 1 | 88.9 | 8.3 | 2.7 |

| | | | |
|---|---|---|---|
| Reaction 2, Set 1 | 89.1 | 7.9 | 3.1 |
| Reaction 1, Set 2 | 89.5 | 7.8 | 2.7 |
| Reaction 2, Set 2 | 89.3 | 8.0 | 2.8 |
| Reaction 1, Set 3 | 89.2 | 8.8 | 2.0 |
| Reaction 2, Set 3 | 88.9 | 9.1 | 2.1 |

[a] Within a given set, reactions are independent replicates initiated simultaneously whereas reactions across different sets are independent replicates performed on separate dates. Given that the ambient flow rates and moisture levels can vary daily, reactions performed across different sets exhibited greater variability compared to those within the same set.
[b,c] $n \geq 2$ replicates.
[d] Individual extension ratio was calculated based on the assumption that all internal sites (5 in total) and terminal sites (2 in total) contributed equally to the extension.
[e] $n = 6$ replicates.

**Table S5. Observed and calculated masses of the hetero-bridged dinucleotides containing tC in a one-pot mixture of 1:1:1 *rC:*araC:*tC. [a]**

| | Observed Mass (Da) | Calculated Mass (Da) |
|---|---|---|
| **rC*tC** | 674.13 | 674.13 |
| **araC*tC** | 663.12 | 663.12 |

[a] Activated mononucleotides (*rC, *araC, and *tC) were mixed in the ratio of 1:1:1. The mixture got spontaneously air-dried. It was then resuspended in LC-MS grade water and immediately submitted for LC-MS analysis.
